# The atlas of RNase H antisense oligonucleotide distribution and activity in the CNS of rodents and non-human primates following central administration

**DOI:** 10.1101/2020.07.31.216721

**Authors:** Paymaan Jafar-nejad, Berit Powers, Armand Soriano, Hien Zhao, Daniel A. Norris, John Matson, Beatrice DeBrosse-Serra, Jamie Watson, Padmakumar Narayanan, Curt Mazur, Holly Kordasiewicz, Eric E. Swayze, Frank Rigo

## Abstract

Antisense oligonucleotides (ASOs) have emerged as a new class of drugs to treat a wide range of diseases, including neurological indications. Spinraza, an ASO that modulates splicing of *SMN2* RNA, has shown profound disease modifying effects in Spinal Muscular Atrophy (SMA) patients, energizing the field to develop ASOs for other neurological diseases. While SMA specifically affects spinal motor neurons, other neurological diseases affect different central nervous system (CNS) regions, neuronal, and non-neuronal cells. Therefore, it is critically important to characterize ASO distribution and activity in all major CNS structures and cell types to have a better understanding of which neurological diseases are amenable to ASO therapy. Here we present for the first time the atlas of ASO distribution and activity in the CNS of mice, rats, and non-human primates (NHP), species commonly used in preclinical therapeutic development. Following central administration of an ASO to rodents, we observe widespread distribution and robust activity throughout the CNS in neurons, oligodendrocytes, astrocytes, and microglia. This is also the case in NHP, despite a larger CNS volume and more complex neuroarchitecture. Our results demonstrate that ASO drugs are well suited for treating a wide range of neurological diseases for which no effective treatments are available.

## INTRODUCTION

Antisense oligonucleotides (ASOs) are chemically modified single-stranded nucleic acids of 16-25 nucleotides in length that bind complementary RNA via Watson-Crick hybridization and can modulate the expression of genes by harnessing a variety of mechanisms (1). To reduce the expression of genes, ASOs can be designed to harness the RNase H1 mechanism (referred to herein as gapmers) (2). RNase H1 recognizes the duplex formed between a DNA-containing ASO and a target RNA and cleaves the RNA, which is subsequently degraded by cellular nucleases. To confer drug-like properties such as enhanced binding affinity and nuclease resistance ASOs are chemically modified. Typically, a gapmer ASO consists of a central stretch of 8-10 DNA nucleotides which is flanked on either side by 2-5 nucleotides with chemical modifications such as 2’-O-methoxyethyl (MOE), constrained ethyl, or locked nucleic acid. Additionally, the phosphodiester internucleotide linkages are replaced with phosphorothioate to further enhance the stability of ASOs to endogenous nucleases (3).

ASOs were first used in the early 90’s to help determine the functions of proteins in the CNS (4–6). However, the first unequivocal proof of ASO activity and disease modification in the CNS of an animal came in 2006 when a MOE-gapmer ASO targeted to SOD1 was administered to the CNS of a rat model of amyotrophic lateral sclerosis (ALS), resulting in SOD1 mRNA reduction and significant slowing of disease progression (7). This work set the stage for an era in which ASOs have been used effectively in a wide range of neurological disease animal models. Over the years, centrally administered ASOs have demonstrated widespread CNS tissue distribution, activity, and substantial phenotypic prevention and/or even reversal in mouse models of Huntington’s (8), SMA (9), Frontotemporal dementia (10), Parkinson’s (11), TDP-43- and C9orf72-linked ALS (12), Alexander disease (13), Pelizaeus-Merzbacher disease (14), Prion disease (15), and multiple forms of Spinocerebellar Ataxia (16–18). Most of these studies were performed with MOE-gapmer ASOs, which is the ASO design that is most commonly being used in the clinic for neurological diseases (19). Pathology in each of these disease models is driven by distinct cell types in distinct regions of the brain or spinal cord, including neurons, astrocytes and oligodendrocytes. This work demonstrates that ASOs can effectively target a wide range of CNS regions and cell types in mice to produce disease-modifying effects.

In larger preclinical species such as rats and NHPs, we have previously demonstrated widespread distribution and activity of MOE-gapmer and uniform MOE ASOs in the CNS following central administration (8,20–23). Using a battery of live imaging techniques and postmortem analyses, MOE-gapmer ASOs were shown to immediately distribute into the cranium following intrathecal (IT) injection in the rat and penetrate all major CNS structures by 6 h post-injection (20,23). Cellular uptake took place between 6-24 hours post-injection and stable regional and cellular distribution was observed from at least 1 to 14 days post-dosing. Reduction of target mRNA and protein was sustained for at least 4-16 weeks post-injection, depending upon the CNS region. Similarly, in NHP live imaging techniques and postmortem tissue analyses were used to demonstrate rapid, widespread, and sustained ASO distribution into the cranium and brain parenchyma, and potent target RNA reduction across the CNS following IT administration via lumbar puncture of a MOE-gapmer ASO targeting the ubiquitously expressed Malat1 non-coding RNA (21).

The pattern of ASO distribution and activity in the CNS of preclinical species is also evident in the human CNS. The first ASO dosed into the human CNS was a MOE-gapmer ASO targeting SOD1, which was administered intrathecally by infusion at low, escalating dose levels in a Phase I study to test its safety and pharmacokinetics in SOD1-linked ALS patients (24). The accumulation of ASO in the CSF was dose-dependent and the concentration of the ASO in the spinal cord tissue obtained at autopsy was consistent with the expected tissue concentration based on preclinical studies in NHPs. In clinical trials for Spinraza, a uniform MOE SMN splicing ASO developed for the treatment of SMA, intrathecal delivery resulted in the accumulation of the ASO in neurons and glial cells in the spinal cord and cortex obtained at autopsy. In addition, there was a robust increase in SMN2 mRNA and SMN protein levels in spinal cord and cortex (25). Spinraza showed substantial efficacy in clinical trials, improving motor function and prolonging survival of SMA infants (26), and became the first centrally administered ASO approved for human use (27). This was a clear demonstration that antisense technology could be harnessed for use in CNS and provided strong motivation for a variety of other neurological diseases to be treated using this platform. Additionally, in clinical trials for a MOE-gapmer ASO targeting Huntingtin (HTT) for Huntington’s disease, dose-responsive ASO concentrations and reductions in mutant HTT levels were observed in the CSF, as a surrogate for HTT level in CNS tissue, and a clear demonstration of pharmacodynamic ASO efficacy in humans (28). In a recent phase 1-2 trial in SOD1-ALS patients, IT administration of Tofersen, a MOE-gapmer ASO targeting SOD1, significantly reduced SOD1 concentrations in CSF and despite the low number of patients enrolled in the study showed a trend towards slowing down the disease progression in the ASO treated patients (29).

Although ASOs have been used successfully in a wide range of mouse models of neurological diseases and in larger preclinical species such as rat and NHPs, the field lacks a comprehensive catalog of ASO distribution and efficacy across all major CNS structures and cell types in preclinical species that would provide guidance for treatment of specific CNS diseases using this drug modality. In this study, we performed a complete characterization of ASO distribution and activity in all the major CNS regions and cell types of rodents and NHPs using a MOE-gapmer ASO targeted to the ubiquitously expressed non-coding and nuclear-retained Malat1 RNA.

## MATERIALS AND METHODS

### ASO preparation

The lyophilized Malat1 ASO (ION-626112, Supplementary table 1) was dissolved in sterile phosphate-buffered saline without calcium or magnesium for experiments in mice and rats, and artificial cerebrospinal fluid (aCSF) for monkeys. The ASO was quantified by UV spectrometry and was sterilized by passage through a 0.2-mm filter before dosing.

### Intracerebroventricular (ICV) administration of ASO in mice

All protocols met ethical standards for animal experimentation and were approved by the Institutional Animal Care and Use Committee of Ionis Pharmaceuticals. Wild-type female mice were obtained from the Jackson Laboratory (C57BL/6J, stock number 000664; Bar Harbor, ME). For ICV bolus injections, mice were placed in a stereotaxic frame and anesthetized with 2% isoflurane by a nose cone fitted into the frame. The scalp and anterior back were then shaved and disinfected. A ~1 cm incision was made in the scalp, and the subcutaneous tissue and periosteum were scraped from the skull with a sterile cotton-tipped applicator. A 10-μl Hamilton microsyringe with a 26 G Huber point removable needle was driven through the skull at 0.3 mm anterior and 1.0 mm lateral to bregma, and was lowered to a depth of 3 mm. Ten microliters of ASO solution was injected a single time into the right lateral ventricle at a rate of 1 μl/s. After 3 min, the needle was slowly withdrawn, and the scalp incision was sutured. The mice were then allowed to recover from the anesthesia in their home cage.

### Intrathecal (IT) administration of ASO in rats

Intrathecal administrations of ASOs in rats were performed, under a protocol approved by the Institutional Animal Care and Use Committee of Ionis Pharmaceuticals. Male Sprague-Dawley rats with body weights between 0.25 and 0.35 kg were obtained from Harlan laboratories (Sprague Dawley, stock number 213M, Livermore, CA). Catheterization of the lumbar intrathecal space was performed as described in Mazur et al. (30). Thirty microliters of either ASO dosing solution or vehicle followed by 40 μl of aCSF vehicle was injected into the subarachnoid intrathecal space via the catheter over approximately 30 s. The animals recovered from anesthesia in their home cage on clean bedding.

### Intrathecal (IT) administration of ASO in nonhuman primates

NHP studies were performed at Charles River Laboratories Montreal ULC and were approved by their Institutional Animal Care and Use Committee. Six adult female cynomolgus monkeys weighing 2.4 ± 0.2 kg were anesthetized with a ketamine/dexmedetomidine/glycopyrrolate cocktail and a reversal agent, atipamezole, was given following the completion of dosing. On Day 1, 14 and 28 artificial cerebrospinal fluid or 25 mg of Malat1 ASO was administered via percutaneous intrathecal injection using a spinal needle at the lumbar level (target L4/L5 space). The dose volume was fixed for all injections at 1 ml and administered over 1 min as a slow bolus. Animals were dosed in lateral recumbency position and remained in a prone position for at least 15 min following dosing prior to the administration of the reversal agent. No significant clinical or biochemical abnormalities were observed in treated animals (data not shown).

### Tissue harvesting

The animals were euthanized 2 weeks after the last dose. Frozen brain and spinal cord tissues were harvested for the determination of ASO concentration and target mRNA expression. Tissues were also immersion fixed in 10% buffered formalin solution for histological processing.

### Real-time RT-PCR

The tissues were homogenized in RLT buffer (Qiagen, Valencia, CA), and 1% (v/v) b-mercaptoethanol. Homogenization was performed for 20 s at 6000 rpm using a FastPrep Automated Homogenizer (MP Biomedicals). Total RNA was then isolated using the RNeasy 96 Kit (Qiagen, Germantown, MD) that included an in-column DNA digestion with 50 U of DNase I (Invitrogen). Real-time RT-PCR was performed with the EXPRESSS One-Step SuperScript qRT-PCR kit (Thermo Fisher Scientific) using gene-specific primers (Supplementary Table 2) (IDT technologies, Coralville, IA). For ASO treated animals the expression level of Malat1 was normalized to that of Gapdh, and this was further normalized to the level in vehicle treated animals.

### Analysis of tissue levels of Malat1 ASO

Samples were weighed, minced, and homogenized with homogenization buffer (20 mM Tris pH 8, 20 mM EDTA, 0.1 M NaCl, 0.5% NP40) and beads (Matrix Green, Bio 101 No. 6040-801) in a 96-well format plate. A calibration curve and quality control samples of known ASO concentrations were prepared in normal homogenized brain and analyzed on the same plate. The ASO was extracted from the tissue, calibration curve, and QC samples via a liquid-liquid extraction using ammonium hydroxide and phenol: chloroform: isoamyl alcohol (25:24:1) (Sigma-Aldrich, St. Louis, MO) and the aqueous layer was then further processed. The ASO concentration in mouse samples was measured by HELISA. (31). Briefly, the aqueous layer was evaporated to dryness and reconstituted in normal K_2_ EDTA human plasma (Bioreclamation). Reconstituted tissue, calibration curve, and QC samples were further diluted in normal human plasma, as necessary, and analyzed via a hybridization ELISA method, on a Molecular Devices Spectramax Gemini XPS (Sunnyvale, CA). The ASO concentration in rat and monkey samples were analyzed by LCMS as previously described (32). The aqueous layer was processed via solid phase extraction with a Strata X plate. Eluates were passed through a protein precipitation plate before drying down under nitrogen at 50°C. Dried samples were reconstituted in 140 μl water containing 100 μM EDTA and were run on an Agilent 6130B single quadrupole LCMS.

### Isolation of CNS cell types from mice

For isolation of cellular subtypes, mice were deeply anesthetized with 3% isoflurane and transcardially perfused with ice-cold PBS (Gibco 14190). A 2-mm piece of cortex ipsilateral to the injection site was dissected and stored at −80 °C until RNA extraction. Remaining cortex from ipsilateral and contralateral sides were dissected and stored in ice-cold Hibernate E (Brain Bits) until being subjected to enzyme digestion to generate single cell suspensions according to the Adult Brain Dissociation Kit (Miltenyi Biotec, 130-107-677). Myelin was removed from the samples using myelin removal beads (Miltenyi Biotec, 130-096-733). Neuron isolation kit (130-098-752), Anti-ACSA-2 microbead kit (130-097-678), CD11b (microglia) microbeads (130-093-634), and Anti-O4 microbeads (130-094-543) were used for magnetic isolation of neurons, astrocytes, microglia, and oligodendrocytes, respectively. Cell viability was determined to be > 90% using Vi-Cell XR cell viability analyzer with trypan blue dye exclusion method (data not shown). Isolated cells were suspended in 300 μl of 1% BME/RLT and frozen at −80°C until RNA extraction.

### Histology

Tissue was fixed and then processed on a mouse brain protocol on a Sakura Tissue Tek tissue processor, after embedding slides were cut at 4 microns, air dried overnight then dried at 60° for 1 hour.

### ASO immunohistochemistry

Slides were deparaffinized and hydrated to water then stained on the Thermo/Labvision Auto-stainer. Sections were incubated in Endogenous Peroxidase Blocker (DAKO, S2003) for 10 Minutes. Followed by proteolytic digestion with Proteinase-K (DAKO S3020) for 1 minute. Additional blocking was done with Background Buster (Innovex, NB306) for 30 minutes. Slides were then incubated with the primary antibody, a polyclonal rabbit anti-ASO antibody (8) at a dilution of 1:10,000 (diluted in 2% bovine serum albumin, 5% donkey serum) for 1 hour. A secondary antibody Donkey anti Rabbit HRP (Jackson, 711-036-152) was applied for 30 minutes. Liquid DAB+ Substrate Chromogen System (Dako, K3468) was used to visualize the ASO antibody. Slides were counterstained with Hematoxylin for 30 seconds then dehydrated, cleared, and coverslipped with Micromount (Leica, 3801731).

### Colorimetric in situ hybridization

In situ hybridization for Malat1 RNA was performed on a Leica Bond RX with a RNAScope custom species-specific probe from Advanced Cell Diagnostics (ACD). The DapB negative control and PPIB positive control probes were used for quality control of the staining procedure. Hybridization was preformed using RNAScope 2.5 LS Reagents Red kit (ACD, 322150). After hybridization the probes were detected with BOND Polymer Refine Red Detection kit (Leica, DS9390). Slides were counterstained with Hematoxylin for 30 seconds then dehydrated, cleared, and coverslipped with Micromount (Leica, 3801731).

### Multiplexed fluorescent in situ hybridization

Double in situ hybridization was performed on a Leica Bond RX with RNAScope custom species-specific probes from ACD. Mouse Malat1 C1 probe was multiplexed with the following mouse C2 probes; Gja1 (astrocytes), Mog (oligodendrocytes), Rbfox3 (neurons) and Tmem119 (microglia) (Supplementary Fig. 1,2). The DapB negative control and PPIB positive control probes were used for quality control of the staining procedure. The RNAScope LS Multiplex Fluorescent Reagent Kit (ACD, 322800) was used during hybridization. Malat1 was detected with a Cy3 and the C2 probes were detected with a Cy5. After labeling slides were rinsed in distilled water, and coverslipped with FluoroGell II with DAPI (EMS, 17985-50) for staining of nuclei. The slides were then imaged on an EVOS2 microscope (Thermo Fisher).

### Data analysis

The ED_50_ was calculated using GraphPad Prism version 6.0 or higher (GraphPad Software, San Diego, CA) after fitting the data using nonlinear regression with normalized response and variable slope. A comparison of the uptake of ASO into CNS tissues between species was accomplished using dose administered, CSF volume, and ASO concentration in CNS tissues. ASO is cleared from the CSF space by presumably 2 mechanisms, 1) bulk flow of ASO to systemic circulation driven by CSF formation/reabsorption, and 2) uptake of ASO into tissues. The initial concentrations in CSF was estimated using the dose administered divided by the CSF volume (Dose/CSF volume) which represents the CNS tissues exposure to ASO and should be a good indicator of uptake into CNS tissues and should be independent of the ASO clearance mechanisms. A linear regression analysis was performed on each species (mouse and rat) and tissue (spinal cord and cortex) combination on a log-log plot of ASO concentration vs. Dose/CSF volume. To account for the difference in the Dose/CSF volume values between species and allow comparisons between the concentrations, a bootstrap analysis was performed to estimate the distribution of predicted ASO concentration values at the Dose/CSF volume value of the observed monkey data (75 mg [total dose] /15 mL CSF volume). A large resampling data set (n=200) was used to estimate the 95% confidence interval across the range of values, while a smaller set (n=24, cortex; n=12, cord) was used to generate a set of data for comparison to observed values. A one-factor ANOVA analysis as well as an all-pairwise t-test multi-comparison test was performed on each set of tissue data.

## RESULTS

### ICV bolus injection of Malat1 ASO results in widespread ASO distribution and target reduction in mice

To evaluate the ASO distribution and activity in the mouse CNS, we administered increasing doses of a Malat1 5-10-5 MOE-gapmer ASO to adult mice by a single ICV bolus injection. Two weeks after the injection, we measured the level of Malat1 RNA and the ASO concentration in a variety of CNS regions (Fig. 1). A dose-dependent reduction of Malat1 RNA was observed in all CNS regions tested and the degree of RNA reduction was correlated with the amount of ASO in the tissue (Fig 1A-H).

**Figure 1.**
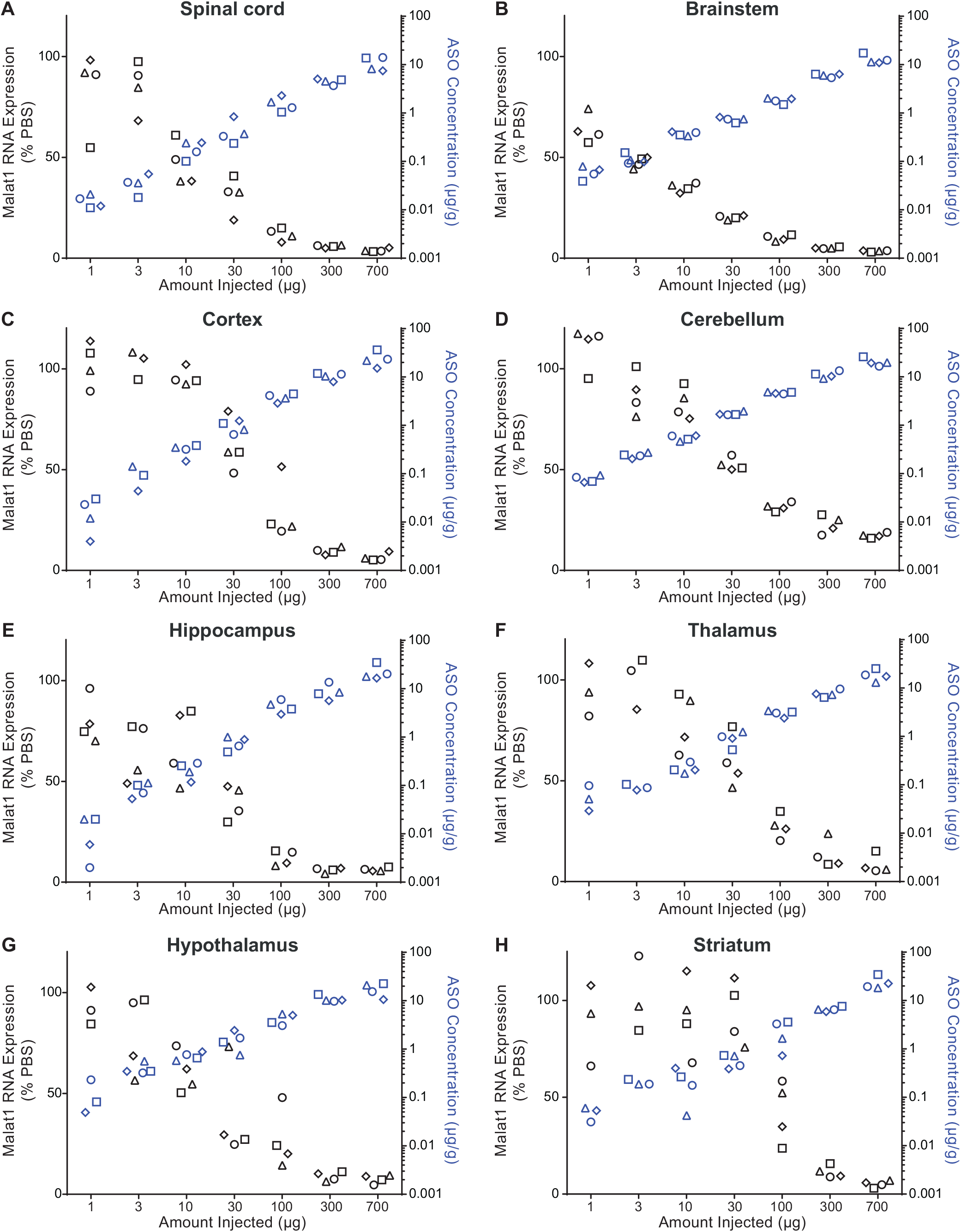
Does responsive Malat1 RNA reduction and ASO accumulation in mouse CNS tissue. Real-time RT-PCR analysis and the MALAT1 ASO tissue concentration (black and blue symbols respectively) are plotted for each animal in each dose for (A) spinal cord, (B) brainstem, (C) cortex, (D) cerebellum, (E) hippocampus, (F) thalamus, (G) hypothalamus, and (H) striatum. MALAT1 ASO concentrations for all CNS regions for the doses 1 μg, 3 μg, 10 μg, and 30 μg were measured by HELISA and for the doses 100 μg, 300 μg, and 700 μg were measured by LC/MS. The left Y axis is for RNA levels which are plotted as % vehicle control and the right Y axis is for ASO concentrations which are plotted as μg ASO per gram of tissue.

Bulk RNA measurements from tissues lack detailed information about the targetability at the sub-regional and cellular level in the CNS. Therefore, to evaluate Malat1 RNA reduction throughout the mouse CNS in more detail, we performed Malat1 in situ hybridization (ISH) on brain and spinal cord of the mice treated with 300 μg of Malat1 ASO. Similar to the qPCR data in Figure 1, ISH data showed that Malat1 RNA was reduced throughout the mouse CNS in all brain regions (Fig. 2A-H). IHC staining with an ASO antibody showed broad distribution throughout the mouse CNS (Fig. 2A-H). In the cerebellum, ISH showed that Malat1 RNA was efficiently reduced in Purkinje cells and the molecular layer of the cerebellum. In contrast, the granular layer of the cerebellum showed much less Malat1 reduction (Fig. 2D; Supplementary Fig. 3). Accordingly, the IHC staining for ASO showed a greater amount of ASO in the molecular and Purkinje cell layers compared to the granular layer. These results demonstrate that ICV administration of the Malat1 ASO results in broad distribution and dose-dependent target reduction in most CNS regions of mice.

**Figure 2.**
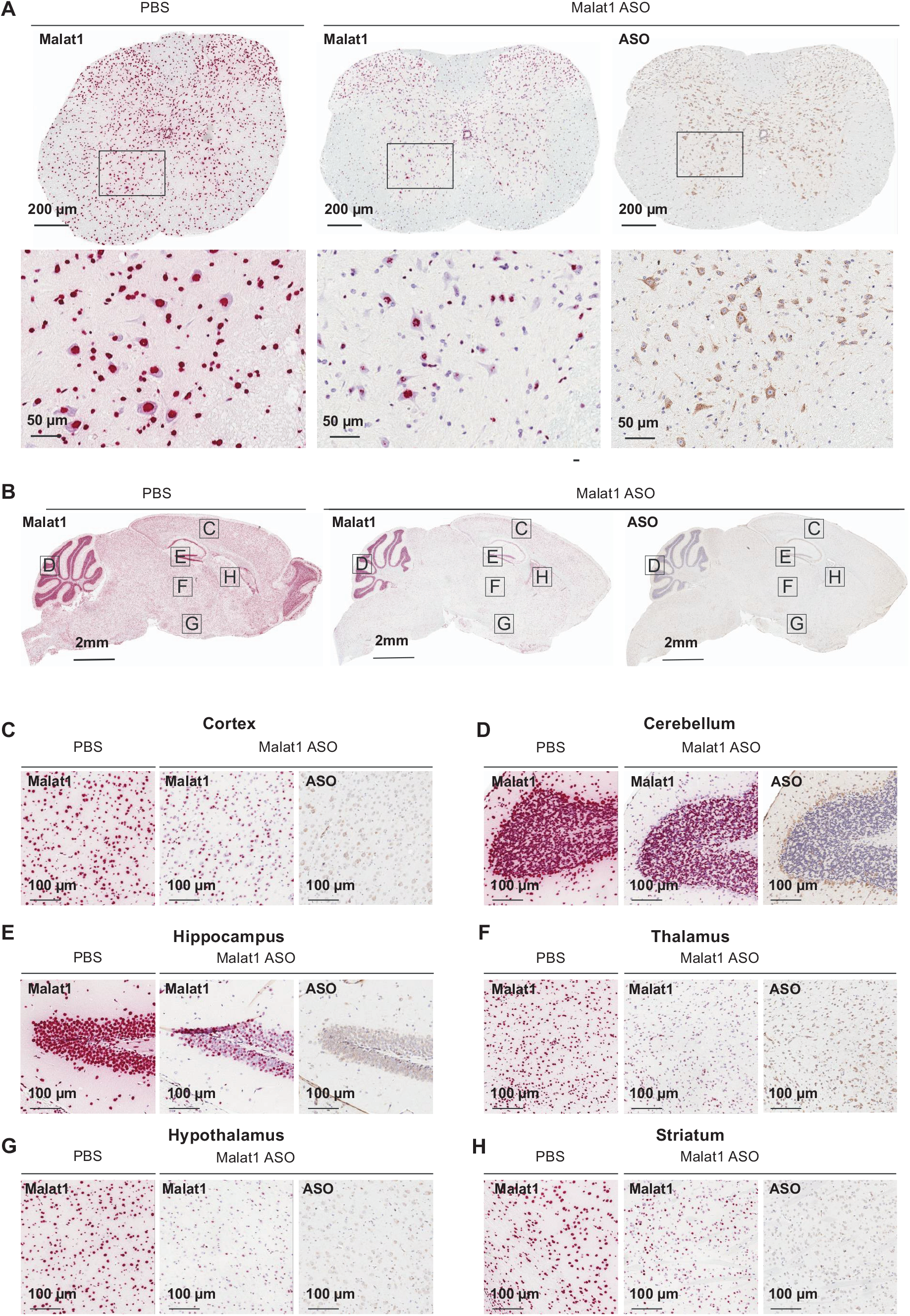
Widespread ASO distribution and Malat1 RNA reduction in mouse CNS. (A) spinal cord, (B) brain, (C) cortex, (D) cerebellum, (E) hippocampus, (F) thalamus, (G) hypothalamus, and (H) striatum were stained for Malat1 RNA and ASO. Staining, treatment, and the scales are indicated. All sections were counter stained with hematoxylin. Mice were treated with 300 μg MALAT1 ASO or vehicle.

### IT bolus injection of Malat1 ASO results in widespread ASO distribution and target reduction in rats

In rats, the CNS and CSF volumes are ~6 times larger than in mice. Therefore, we asked if the distribution and activity of the Malat1 ASO was as robust in rats as was observed in mice. Moreover, the larger size of the rats compared to mice provided the opportunity to perform IT delivery to mimic a clinically relevant route of administration. Increasing doses of the Malat1 ASO were administered to adult rats by a single IT bolus injection. After two weeks we dissected a variety of CNS regions and measured the levels of Malat1 RNA and the ASO tissue concentration (Fig. 3). As we demonstrated in mice, dose-dependent reduction of Malat1 RNA was observed in all CNS regions tested and the degree of RNA reduction was correlated with the amount of ASO in the tissue (Fig. 3A-M). The ASO concentration in the striatum even at the highest dose tested was lower than in other CNS regions and therefore only ~50% Malat1 RNA reduction was observed (Fig. 3M). ISH performed in tissue sections for Malat1 RNA corroborated the qPCR results (Fig. 4A-H). As in mice, IHC staining with an ASO antibody showed broad ASO distribution and ISH staining showed robust reduction of Malat1 RNA. Again, Malat1 ISH in the cerebellum revealed that unlike the molecular and Purkinje cell layers, which were very sensitive to ASO treatment, the granular layer showed modest RNA reduction (Fig. 4D). This provides an explanation for why Malat1 reduction in whole tissue cerebellar extract was not as robust as for most of the other regions (Fig. 3I). These results demonstrate that IT delivery of the Malat1 ASO results in broad distribution and dose-dependent target reduction in most CNS regions of rats.

**Figure 3.**
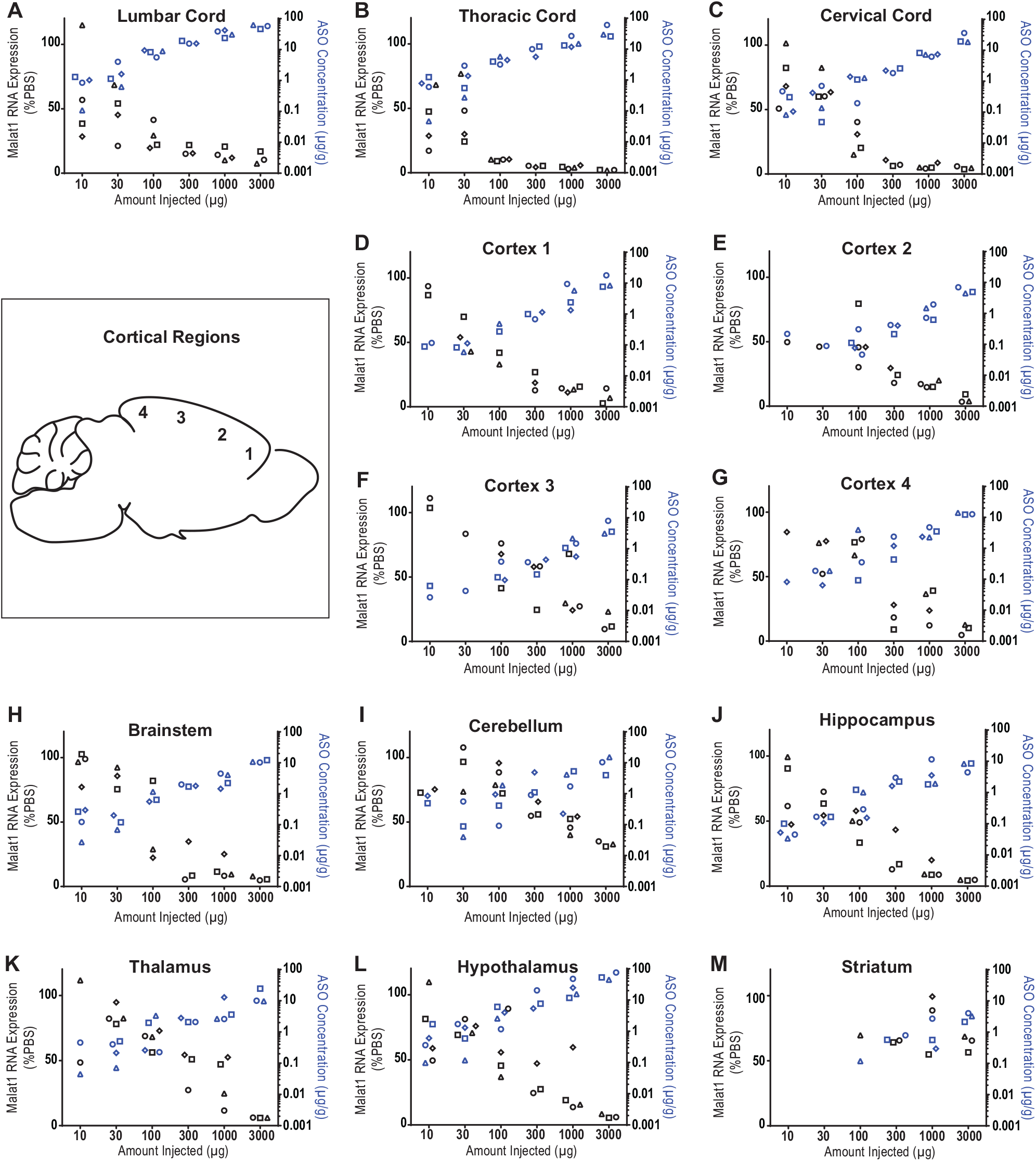
Does responsive Malat1 RNA reduction and ASO accumulation in rat CNS tissue. Real-time RT-PCR analysis and the MALAT1 ASO tissue concentration (black and blue symbols respectively) are plotted for each animal in each dose for spinal cord (A) lumbar, (B) thoracic, and (C) cervical (D-G) cortex regions 1-4, (H) brainstem, (I) cerebellum, (J) hippocampus, (K) thalamus, (L) hypothalamus, and (M) striatum. MALAT1 ASO concentrations for all CNS regions for all doses were measured by LC/MS. The left Y axis is for RNA levels which are plotted as % vehicle control and the right Y axis is for ASO concentrations which are plotted as μg ASO per gram of tissue.

**Figure 4.**
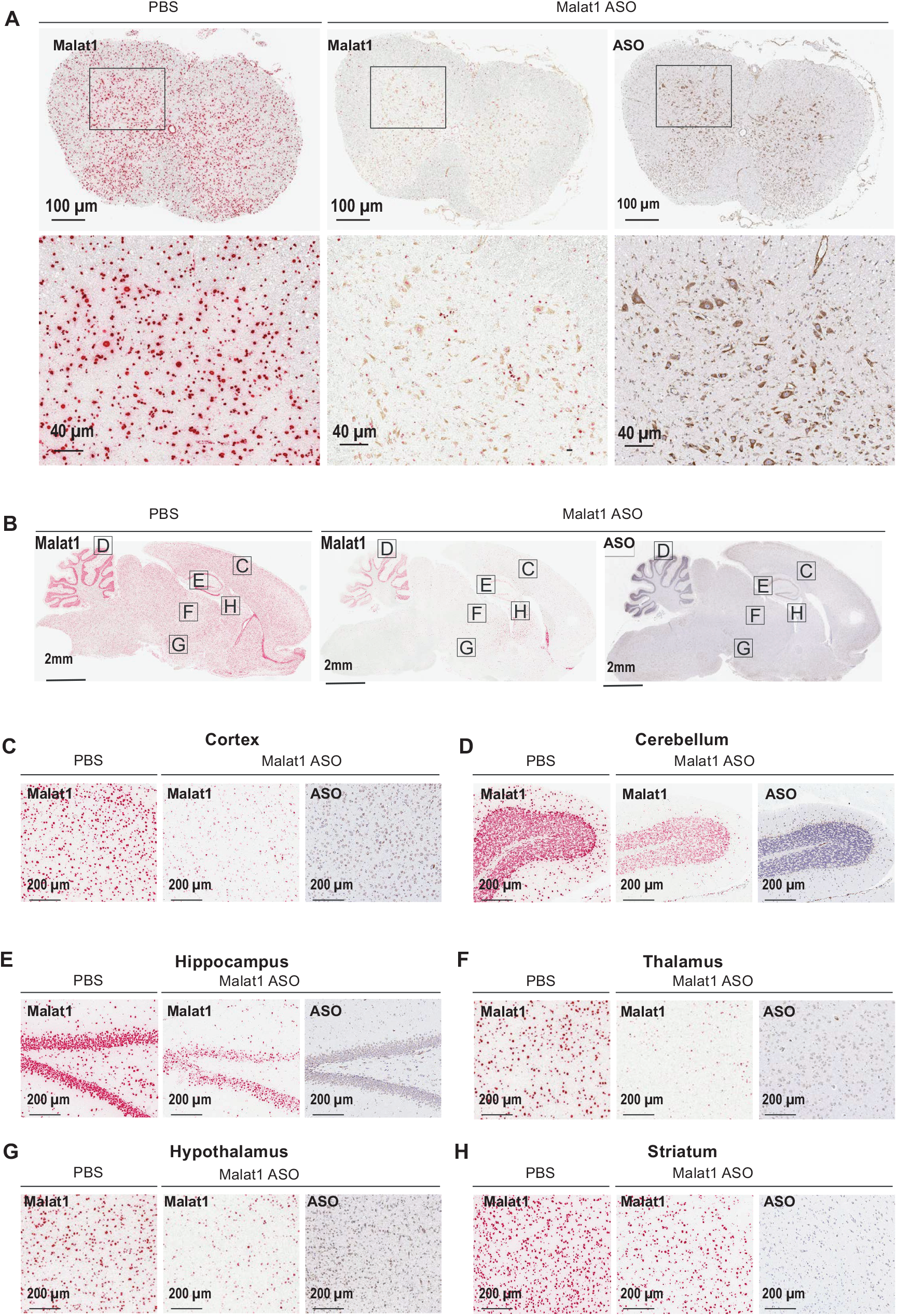
Widespread ASO distribution and Malat1 RNA reduction in rat CNS. (A) spinal cord, (B) brain, (C) cortex, (D) cerebellum, (E) hippocampus, (F) thalamus, (G) hypothalamus, and (H) striatum were stained for Malat1 RNA and ASO. Staining, treatment, and the scales are indicated. All sections were counter stained with hematoxylin. Rats were treated with 1000 μg MALAT1 ASO or vehicle

### IT bolus injection of Malat1 ASO results in widespread ASO distribution and target reduction in NHPs

The CNS and CSF volumes of NHPs are ~60 times larger than those of rats and NHPs are one of the large animal species frequently used for pre-clinical safety evaluation of drugs prior to clinical trials in humans. To evaluate the ASO distribution and Malat1 RNA reduction in the NHP CNS, we administered the Malat1 ASO to adult monkeys by repeated IT bolus injections. We observed broad ASO distribution and Malat1 RNA reduction throughout the CNS (Fig. 5). The ASO concentration in the spinal cord, both proximal and distal to the injection site were high (25 and 13 μg/g, respectively). The highest ASO tissue concentrations were observed in the cortex, with highly consistent concentrations in all cortical areas from frontal cortex to occipital cortex. ASO tissue concentrations in brain regions such as pons, medulla, hippocampus, thalamus, amygdala, and cerebellar cortex and were consistent with each other and similar to the spinal cord levels. ASO tissue concentrations in the globus pallidus, caudate, and putamen were lower compared with other brain regions. The reduction of Malat1 RNA in CNS tissues was in agreement with the ASO tissue concentration, with most regions showing 80-90% target reduction (Fig. 5A). Interestingly, one of the monkeys consistently showed lower ASO concentration across all the tissues, likely due to a suboptimal IT injection, which resulted in less Malat1 reduction. Due to the small size of several regions in the brain it was not possible to perform both ASO quantitation and RNA expression analyses. However, Malat1 RNA was strongly reduced in most regions analyzed (Fig. 5B). Broad ASO distribution and activity in different brain structures offers the opportunity to develop ASO drugs for many CNS diseases.

**Figure 5.**
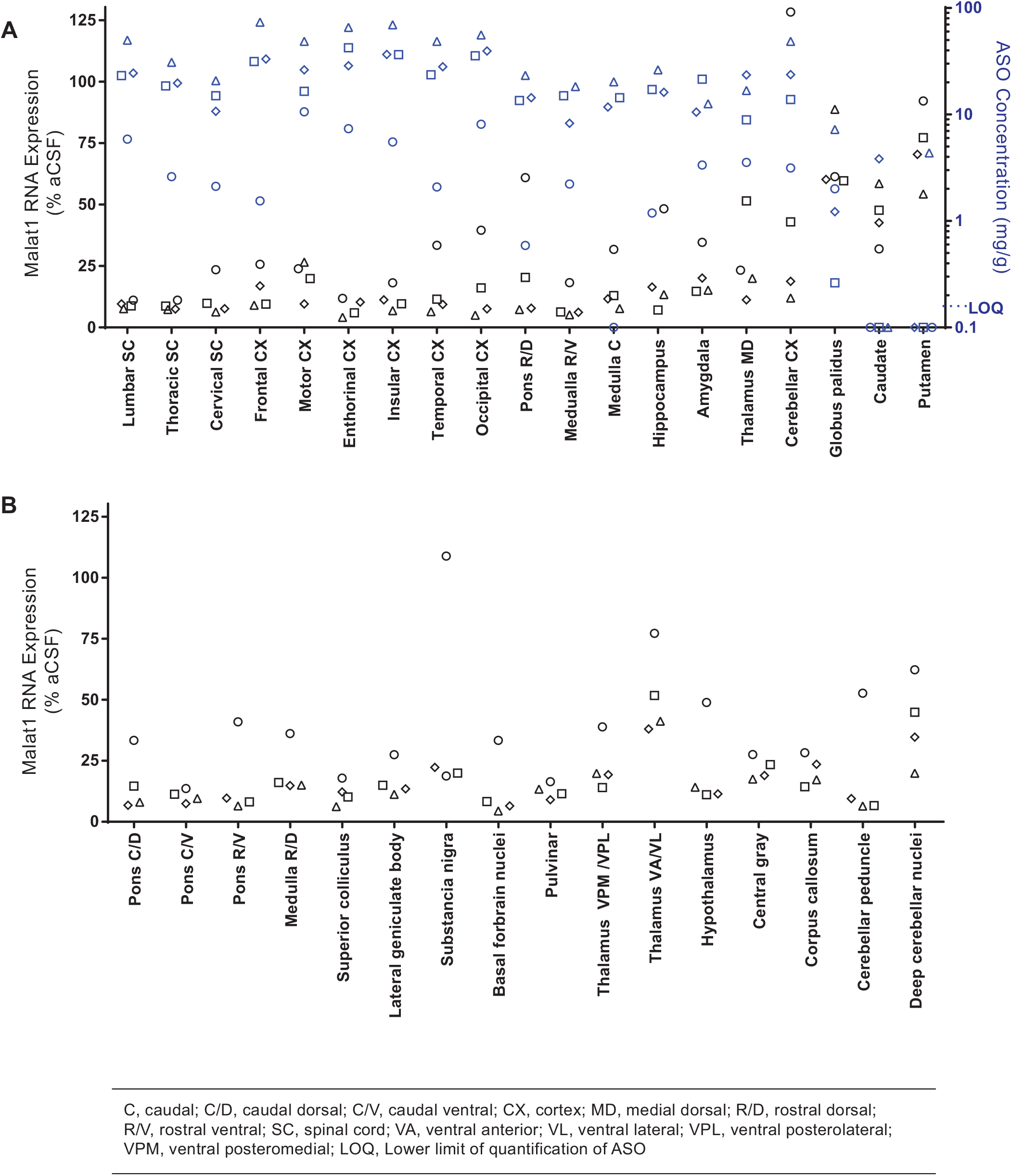
Widespread ASO accumulation and Malat1 RNA reduction in the NHP CNS. (A) Real-time RT-PCR analysis and the MALAT1 ASO tissue concentration (black and blue symbols respectively) are plotted for each animal in several CNS regions. MALAT1 ASO concentrations for all CNS regions for all doses were measured by LC/MS. The left Y axis is for RNA levels which are plotted as % vehicle control and the right Y axis is for ASO concentrations which are plotted as μg ASO per gram of tissue. (B) Real-time RT-PCR analysis for additional CNS regions that were too small to be sampled for ASO concentration analysis. RNA levels are plotted for each animal as % vehicle control.

To assess the ASO distribution within CNS structures in more detail we performed IHC staining with an ASO antibody (Fig. 6–7). The ASO distribution assessed by IHC was in agreement with the ASO concentrations measured in different brain structures. The Malat1 ISH studies also allow for visualization of target reduction in individual cells within a brain structure. Despite the globus pallidus, caudate, and putamen showing the least whole tissue ASO accumulation, ISH staining in these regions reveals clear Malat1 RNA reduction in ASO treated animals at the cellular level (Fig. 7A-C). Trigeminal ganglion and dorsal root ganglia (DRG) at cervical, thoracic, and lumbar levels showed accumulation of ASO accompanied with Malat1 RNA reduction visualized by in situ (Fig. 7H-K). These results show that IT delivery of the Malat1 ASO results in broad distribution and target reduction in most CNS regions of NHPs.

**Figure 6.**
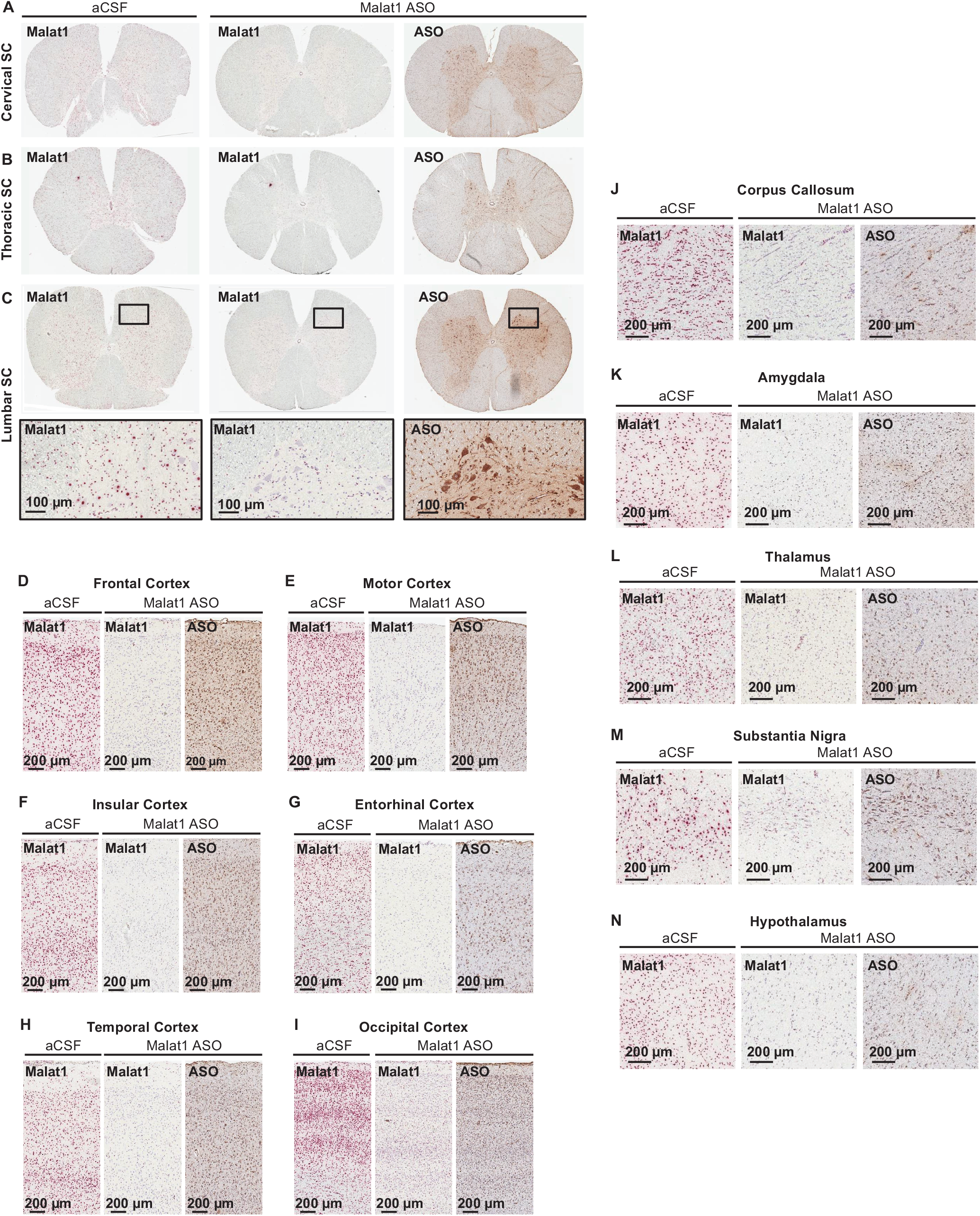
Widespread ASO distribution and Malat1 RNA reduction in the NHP CNS. (A) cervical, (B) thoracic, and (C) lumbar spinal cord, (D) frontal, (E) motor, (F) insular, (G) entorhinal, (H) temporal, and (I) occipital cortex, (J) corpus callosum, (K) amygdala, (L) thalamus, (M) substantia nigra, (N) hypothalamus

**Figure 7.**
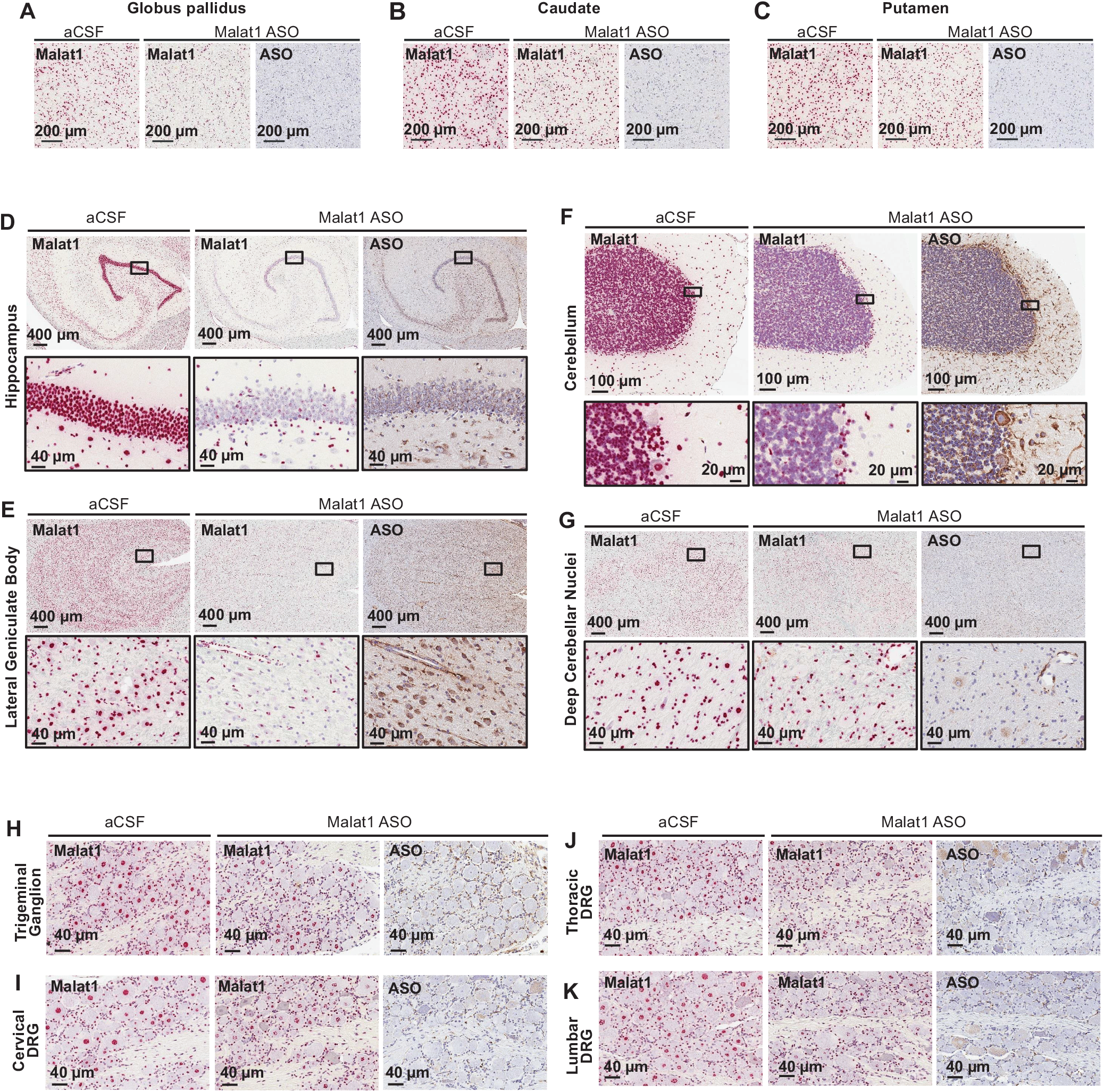
Widespread ASO distribution and Malat1 RNA reduction in the NHP CNS. (A) globus pallidus, (B) caudate, (C) putamen, (D) hippocampus, (E) lateral geniculate body, (F) cerebellum, (G) deep cerebellar nuclei, (H) trigeminal ganglion, (I) cervical, (J) thoracic, and (K) lumbar dorsal root ganglia are stained for Malat1 RNA and ASO. Staining, treatment, and the scales are indicated. All sections were counter stained with hematoxylin. DRG, dorsal root ganglia.

### Central administration of Malat1 ASO results in robust target reduction in all the major CNS cell types

CNS is a complex tissue that consists of four major cell types. Neurons, oligodendrocytes, astrocytes, and microglia play important roles in the pathogenesis of different neurological diseases. We have shown that ASO distributes well throughout the CNS of the three species we examined (Fig. 2, 4, 6, and 7). However, it is equally important to determine how well ASOs can target the different cell types in the CNS. Initially, we noticed that ASO distribution and Malat1 RNA reduction were evident in both white and grey matter regions, and a wide range of cell types based on their location and morphology (Fig 2A-H). To confirm the activity of the Malat1 ASO in the four major cell types of the CNS across the three species, we performed double labelling in situ hybridization using a probe for Malat1 RNA and a probe for each cell type of interest. To label neurons, oligodendrocytes, microglia and astrocytes we used species specific in situ probes for Rbfox3, Mog, Tmem119, and Gja1, respectively (See supplementary methods). Malat1 was robustly reduced in all four cell types of the CNS in all three of the species evaluated (Fig 8A-C).

**Figure 8.**
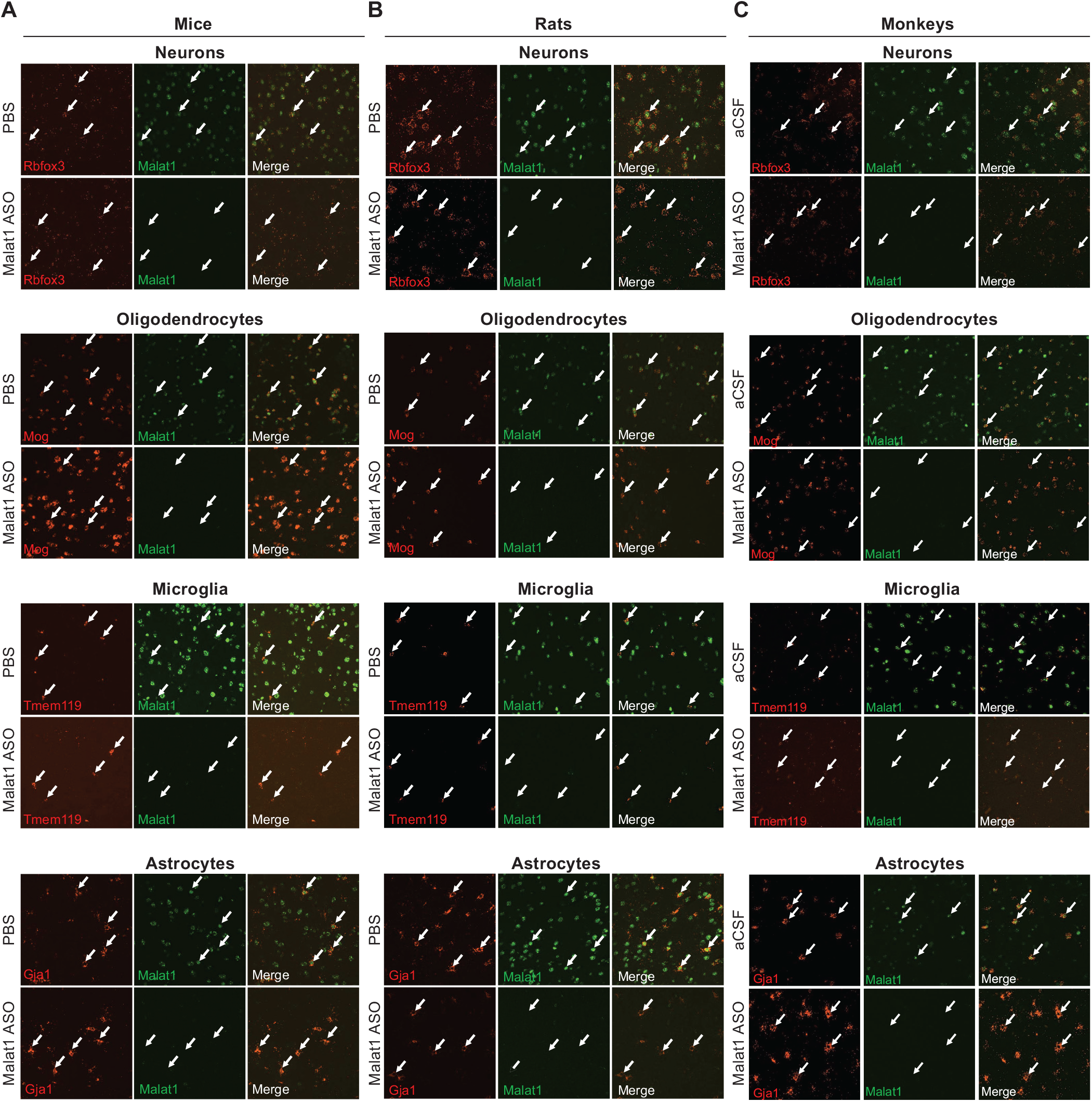
The Malat1 ASO targets all major cell types in the CNS. Colocalization of Malat1 with Rbfox3 in neurons, Mog in oligodendrocytes, Tmem119 in microglia, and Gj1 in astrocytes in cortex of (A) mouse, (B) rat, and (C) NHP. Arrows indicate four selected cells in each field. Mice were treated with 300 μg MALAT1 ASO or vehicle. Rats were treated with 1000 μg MALAT1 ASO or vehicle. NHP were treated with 3 doses of 25 mg MALAT1 ASO or vehicle.

To quantitatively assess Malat1 RNA reduction in the four major CNS cell types, we isolated them from cortex using a magnetic cell isolation and measured the levels of Malat1 RNA by real-time RT-PCR after administering the Malat1 ASO at increasing doses to adult mice by a single ICV bolus injection. As expected, Malat1 RNA was reduced dose-dependently in neurons with an ED_50_ of 206 μg. Surprisingly, microglia (ED_50_ of 71 μg), oligodendrocytes (ED_50_ of 34 μg) and astrocytes (ED_50_ of 17 μg) were more sensitive to the Malat1 ASO compared to neurons (Fig. 9). These data demonstrate that all the major CNS cell types can be robustly targeted by an ASO.

**Figure 9.**
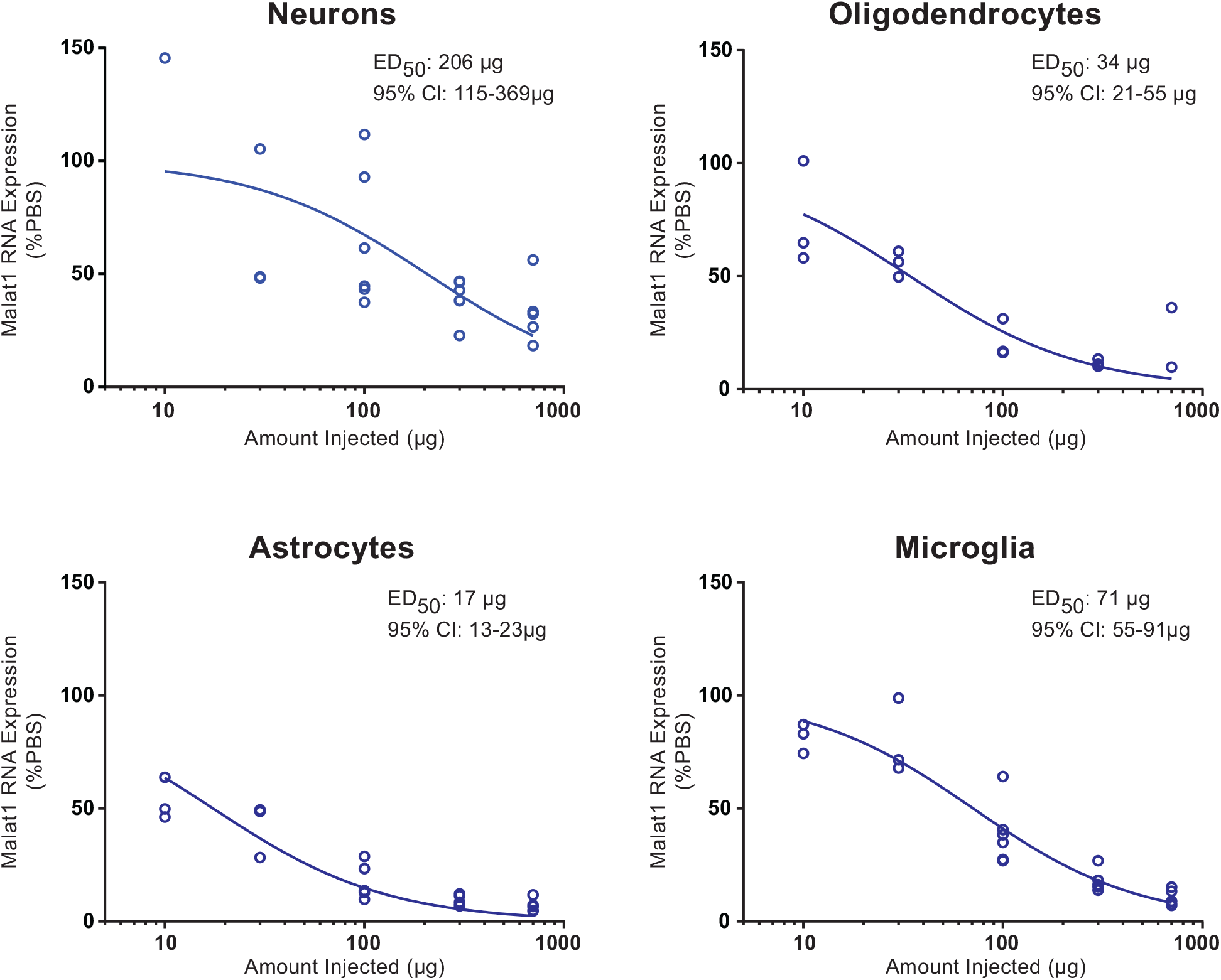
Dose-dependent Malat1 RNA reduction in neurons, oligodendrocytes, astrocytes, and microglia isolated from mouse cortex. Real-time RT-PCR analysis for Malat1 RNA are plotted for individual animals in different cell types. Injected doses are indicated on the X axis.

### CSF volume scaling across species to predict ASO tissue concentration

Since we evaluated ASO concentrations in the CNS tissues of three species with different CNS volumes, we asked if the ASO tissue concentrations scaled accordingly between species. To determine if this was the case, we plotted the ASO dose normalized to CSF volume of each species (24,32) versus the measured ASO tissue concentrations. We predicted the tissue concentration for monkey based on a linear regression analysis using the measured ASO tissue concentrations in either mouse or rat spinal cord and compared this to the measured ASO concentrations in ASO treated monkeys. When the ASO dose was normalized to the CSF volume, the ASO tissue concentration in the mouse spinal cord was significantly less than the ASO tissue concentration in the rat spinal cord (p ≤ 0.0001) (Figure 10A, Supplementary table 3 and 7). The predicted monkey ASO tissue concentration from the mouse spinal cord data was significantly lower than the measured tissue concentration in monkey (p ≤ 0.0001) (Figure 10A, Supplementary table 5 and 8). On the contrary, the predicted monkey ASO tissue concentration from the rat spinal cord data was not significantly different from the measured tissue concentration in monkey (p ≤ 0.8622) (Figure 10A, Supplementary table 5 and 7). Comparing ASO tissue concentrations in the cortex of mouse and rat showed a small but significant difference (p ≤ 0.002), with higher tissue concentration in mouse. However, the ASO tissue concentration measured in monkey was significantly higher than the predicted ASO tissue concentration from both the mouse and rat data (p ≤ 0.0001) (Figure 10B, Supplementary table 4, 6, and 8).

**Figure 10.**
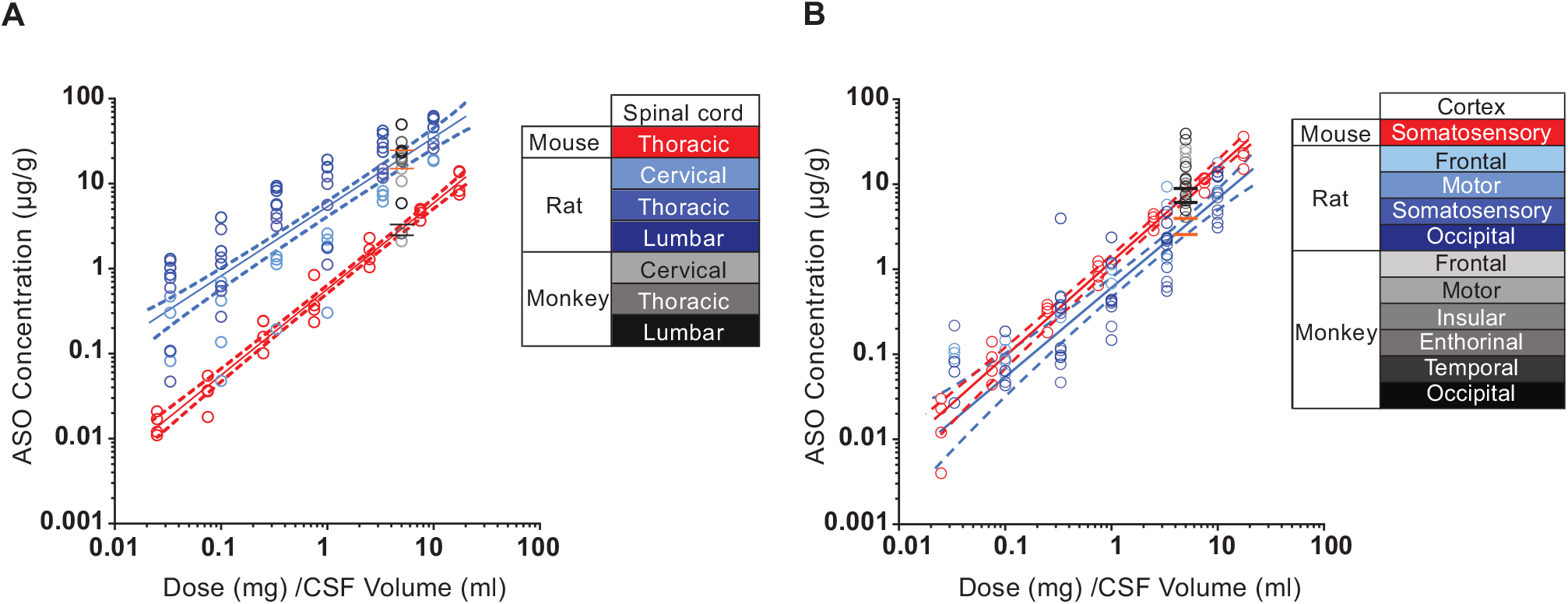
Comparison of ASO tissue concentration after dose normalized to CSF volume in mice, rats, and monkeys in A) spinal cord and B) cortex. The CSF volumes used for mouse, rat and monkey were 0.04 ml, 0.3 ml, and 15 ml, respectively. Data for mice, rats, and monkeys are shown in red, blue and grey symbols, respectively. Different tissues sampled from rats and monkeys are depicted in shades of blue and grey, respectively. The best fit regression is shown as a in solid line (red for mice and blue for rats) and the 95% confidence intervals are shown as dashed lines. The predicted NHP tissue concentrations simulated form boot strap regression for mouse (between the black lines) or rat (between the orange lines) are shown.

## DISCUSSION

Here we demonstrate that central administration of a MOE-gapmer ASO results in widespread distribution and potent target RNA reduction throughout the CNS of rodents and NHPs. The four major CNS cell types, neurons, microglia, astrocytes and oligodendrocytes are all robustly targeted with this ASO.

Across all neurological diseases many different CNS regions and cell types have been shown to play a role. For example, selective spinal motor neuron degeneration occurs in SMA (33), motor neuron loss in both cortex and spinal cord occurs in ALS (34), selective degeneration of substantia nigra pars compacta dopaminergic neurons occurs in Parkinson’s disease (35), degeneration of neurons in the frontal cortex and hippocampus occurs in Alzheimer’s disease (36), and Purkinje cells selectively degenerate in a variety of cerebellar ataxias (37). Since our results demonstrate that there is widespread distribution and activity in most CNS regions and the major cells types, we believe that ASO drugs are well suited for treating a wide range of neurological diseases for which no effective treatments are available.

Some CNS regions and neuron subtypes such as striatum in mice and rats, the caudate nucleus, putamen, and globus pallidus in NHP, and the granule cells in the cerebellum showed less ASO accumulation and Malat1 RNA knock down. For diseases affecting these CNS regions and neuronal subtypes, higher doses of ASO are required to achieve the desired level of mRNA suppression. In the future, advancements in delivery of ASO via ligand-conjugated antisense (LICA) could be used to more robustly target CNS cell types and regions which are less sensitive to unconjugated ASO. This strategy has worked remarkably well to specifically deliver ASOs to hepatocytes using N-acetylgalactosamine (GalNAc) (38) as a ligand or to specifically delivery of ASOs to pancreatic beta cells using glucagon-like peptide 1 (GLP1) as a ligand(39,40). In addition, delivery across the brain-blood barrier (BBB) may offer an opportunity to achieve even broader ASO distribution and activity in the CNS.

While neuronal dysfunction is frequently observed in many neurological diseases, dysfunction of non-neuronal cells such as microglia, oligodendrocytes, and astrocytes can also modulate or drive disease (41,42). Importantly, here we observed robust activity in neurons as well as in the three major glial cell types in rodents and NHPs. For each of the major CNS cell types, though, there are subtypes, and more work is required to fully characterize how sensitive they are to ASOs. In the future, single-cell RNA-seq will help to augment and expand our quantitative assessments of ASO activity in cellular subtypes.

It is gratifying to observe widespread ASO distribution and activity in small animals such as mice to larger animals such as NHPs after central administration. Recently, it was shown that a divalent siRNA has broad distribution and activity in the CNS of NHPs including neurons and GFAP positive astrocytes (43). In contrast to ASOs and divalent siRNAs, intrathecal delivery of other drug modalities such as antibodies, proteins, or peptides results in a large rostral to caudal gradients and limited accumulation in tissue (44,45). Viral delivery of gene therapy achieves good CNS distribution but does not effectively target non-neuronal cells (46). Recently, we showed that following IT bolus injection in rats, ASO distributes rostrally in the CSF along the neuroaxis and rapidly adheres to pia and arterial walls, consistent with perivascular entry into the CNS (20). Over time, ASO accumulates in all major CNS cell types presumably by receptor mediated endocytosis (47). However, more work is needed to better understand the molecular mechanisms responsible for ASO distribution into the tissue and uptake of the ASO into the various CNS cell types.

Since we evaluated the concentration of ASO in the CNS tissues of three species with different CNS volumes, we asked if the ASO tissue concentrations scaled accordingly between species. Our data show that, the ASO tissue concentration in the rat spinal cord following IT delivery of ASO can predict monkey ASO tissue concentration for a given dose when normalized to CSF volume. However, ASO tissue concentrations measured in mouse spinal cord predicted a much lower ASO tissue concentration than the actual measurements in monkey spinal cord. Interestingly, the ASO concentrations in cortex from both mice and rats underestimated monkey ASO tissue concentrations when the dose normalized by CSF volume. ASO is cleared through bulk flow with CSF reabsorption as well as uptake to tissues. The CSF formation and reabsorption rates differ between species (48) which may explain the observed difference in uptake to CNS tissues. Alternatively, differences in anatomical features, and dosing routes (ICV versus IT) could play a role in the differences in uptake observed.

Supplementary Data are available at NAR online.

## ACKNOWLEDGEMENT

The authors thank the vivarium staff, oligo synthesis group, and histology core staff at Ionis Pharmaceuticals Inc, for their technical support.

## FUNDING

Not applicable.

## CONFLICT OF INTEREST

All authors are paid employees of Ionis pharmaceuticals.

## Supplementary data

### Supplementary methods

#### Development of multiplexed fluorescent in situ methods to evaluate ASO activity in different CNS cell types

In order to develop methods to examine cell-type specific RNA reduction in tissue sections, we had to first determine high quality cell type in situ stains to identify neurons, oligodendrocytes, astrocytes and microglia. We started by considering the mRNA for standard cellular antibody stains for the different cell types. We chose NeuN (Rbfox3) for neurons, MOG (Mog) for oligodendrocytes, GFAP (Gfap) for astrocytes, and IBA1 (Aif1) for microglia as immunohistochemistry for these markers has been used for decades to identify the different cell types. We worked with Advanced Cellular Dynamics (ACD) and their RNAScope technology to develop in situ oligonucleotide probes for these targets in the rat. We then euthanized a normal male Sprague-Dawley rat, harvested the brain and spinal cord and processed them for histology. We sent the sections to the Pharma Services division of ACD to test the staining of these probes on normal tissue so that we could evaluate their use as cell type markers. We found that Rbfox3 was a good marker to identify neurons and Mog proved to be a good marker to identify oligodendrocytes (Supplementary Figure 1) so these markers were used in the ASO experiments. In situ staining for astrocytes with Gfap showed a heterogenous staining pattern where not every astrocyte in every structure was identified (data not shown). The in situ staining for Aif1 resulted in staining in what looks like the ends of the fine ramifications of the microglia producing a cloud of staining around an unidentified nucleus (data not shown), and this staining pattern would not work to identify the microglia for analysis of Malat1 RNA expression.

We went on to search for another marker that would identify a larger proportion of the astrocytes in the CNS. We used the website brainrnaseq.org to identify candidate genes that had high, specific astrocyte expression when compared to other cell types. These were ranked based on expression level and astrocyte specificity. Leading candidates were then evaluated on the Allen Brain Atlas website (brain-map.org) for uniformity of expression across the brain. Two candidates were identified, Ndrg2 and Gja1, that had good specificity, high expression level and homogenous expression across the brain. Probes for both genes were made by ACD and tried on the naïve rat tissues. The in-situ staining for Ndrg2 proved to be too homogenous across the tissue and did not delineate the cell bodies well (data not shown). Whereas, the in situ stain for Gja1 delineated the cell bodies of the astrocytes very well and was selected for use in the ASO experiments (Supplementary Figure 2A,B). We also searched for a more suitable in situ stain for microglia using a similar approach. Initially we searched on the Barres website for highly expressed microglia specific markers. Bennett et al. identified Tmem119 as a good marker for most microglia (49) and this was one of our top candidates in the expression/specificity search. An in-situ probe for Tmem119 was made by ACD and used to stain normal rat CNS tissues. We found that in situ staining for Tmem119 clearly delineated the cell bodies of the microglia and was used in the ASO experiments (Supplementary Figure 2C,D).

## Supplementary figures

**Supplementary Figure 1.**
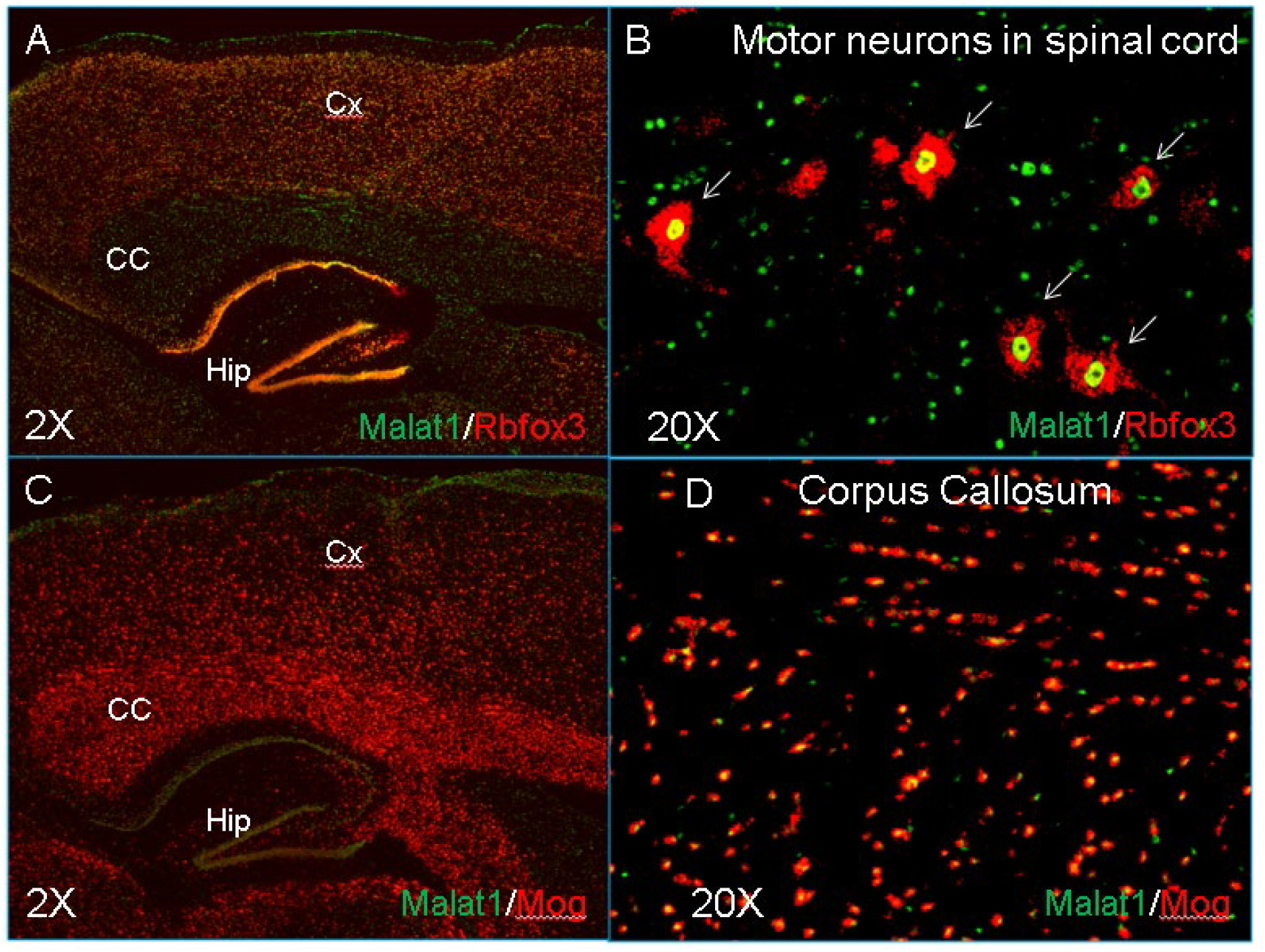
Staining pattern for multiplexed in situ cell type markers (red) Rbfox3 for neurons and Mog for oligodendrocytes and Malat1 (green) in the rat. Cx: cortex, CC: corpus callosum, Hip: hippocampus. White arrows in image B indicate probable motor neurons of the spinal cord. Images A and C are 2X magnification, images B and D are 20X magnification.

**Supplementary Figure 2:**
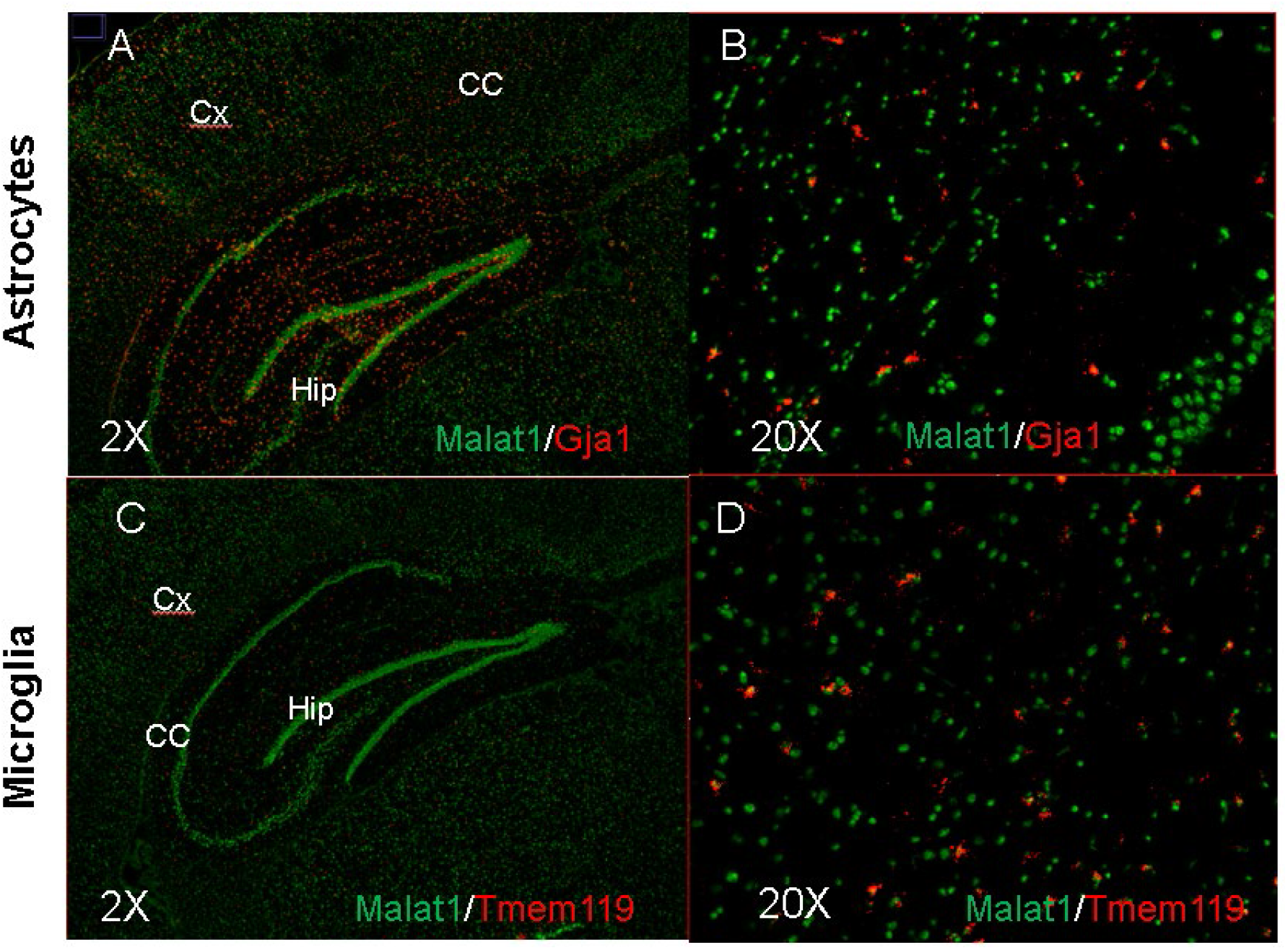
staining pattern for multiplexed in situ cell type markers (red) Gja1 for astrocytes and Tmem119 for microglia and Malat1 (green) in the rat. Cx: cortex, CC: corpus callosum, Hip: hippocampus. Images A and C are 2X magnification, images B and D are 20X magnification.

**Supplementary Figure 3.**
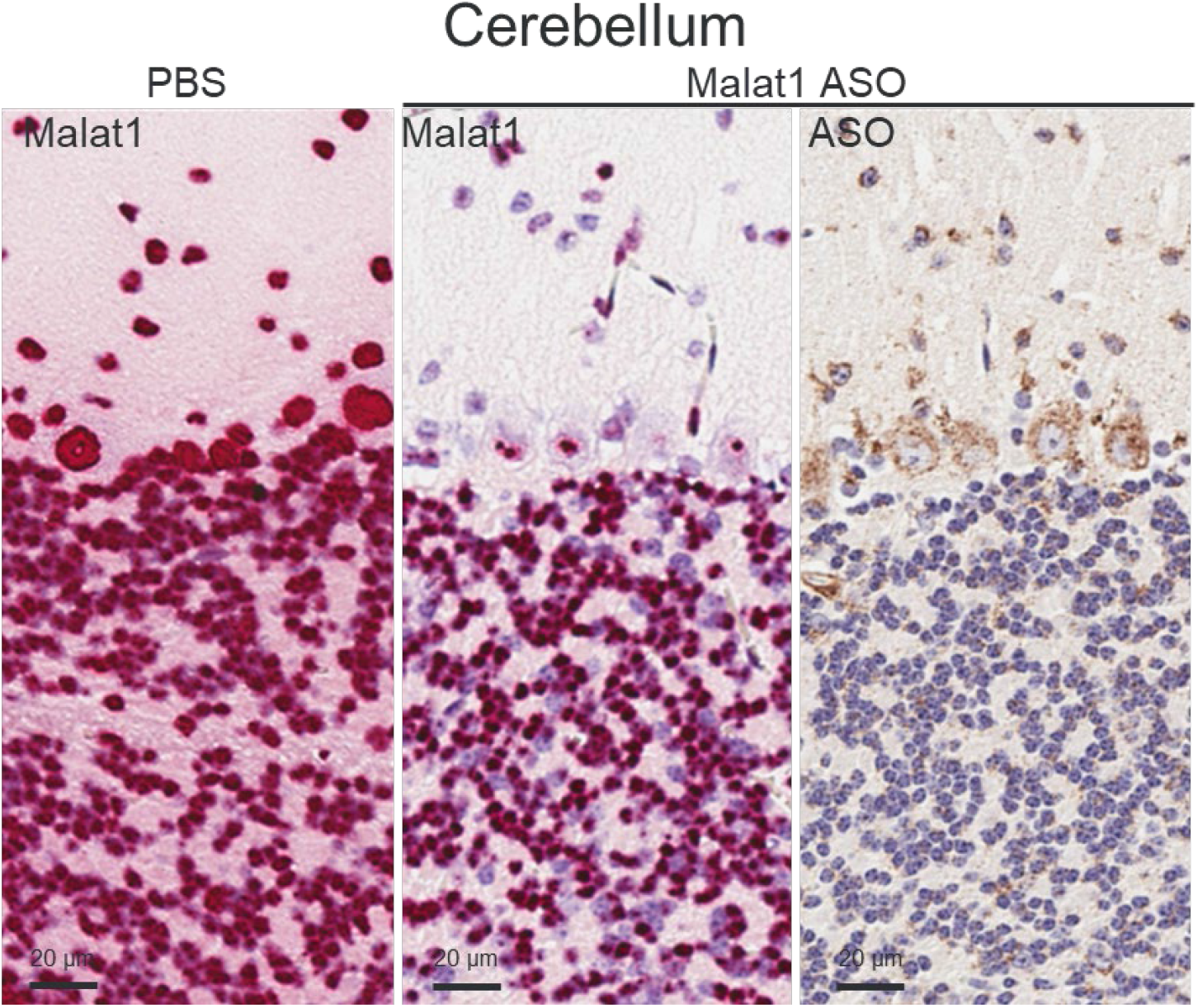
In situ hybridization of Malat1 and ASO immunohistochemistry in mouse cerebellar cortex. Staining, treatment, and the scales are indicated. All sections were counterstained with hematoxylin.

**Supplementary table 1.**
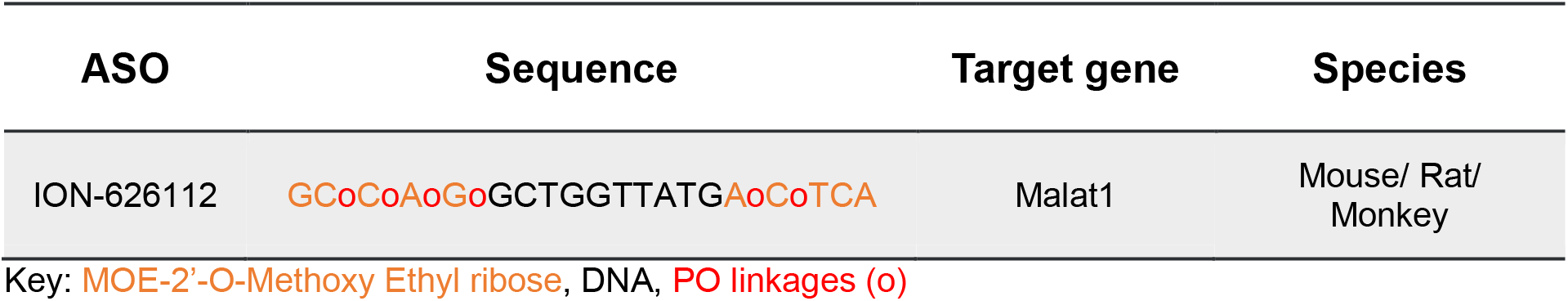
Sequences of antisense oligonucleotide (ASO) used in the study.

**Supplementary table 2.**
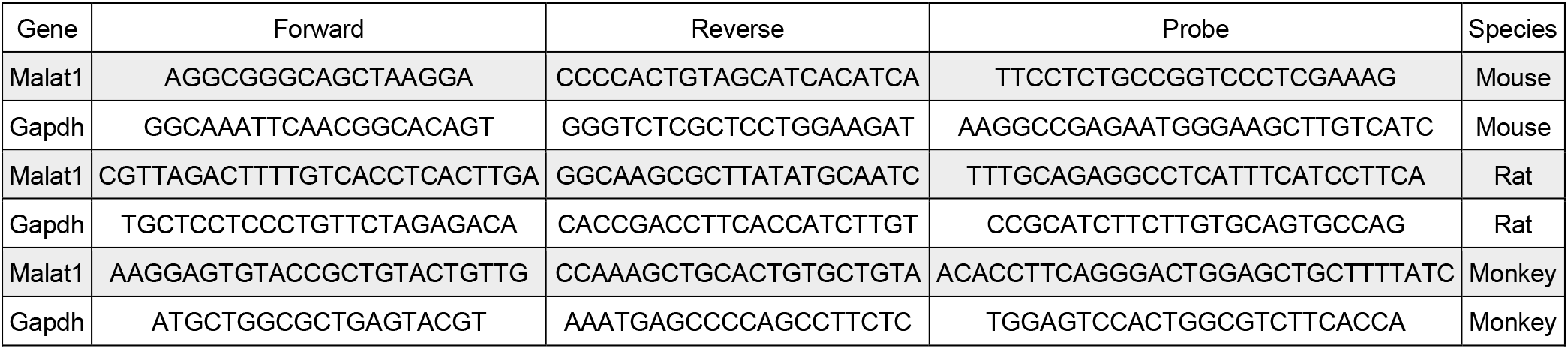
Primer sequences used in the study.

**Supplementary table 3.** Spinal cord ASO tissue concentrations used in figure 10A.

| Species | Tissue (spinal cord) | Dose (mg) | CSF volume | Dose (mg) / CSF volume | ASO concentration (µg) |  |  |  |
| --- | --- | --- | --- | --- | --- | --- | --- | --- |
|  |  |  |  |  | Animal #1 | Animal #2 | Animal #3 | Animal #4 |
| Mouse | Thoracic | 0.001 | 0.04 | 0.025 | 0.02 | 0.01 | 0.02 | 0.01 |
| Mouse | Thoracic | 0.003 | 0.04 | 0.075 | 0.04 | 0.02 | 0.04 | 0.06 |
| Mouse | Thoracic | 0.01 | 0.04 | 0.25 | 0.16 | 0.10 | 0.24 | 0.24 |
| Mouse | Thoracic | 0.03 | 0.04 | 0.75 | 0.33 | 0.24 | 0.37 | 0.84 |
| Mouse | Thoracic | 0.1 | 0.04 | 2.5 | 1.29 | 1.04 | 1.69 | 2.30 |
| Mouse | Thoracic | 0.3 | 0.04 | 7.5 | 3.67 | 4.86 | 4.55 | 5.05 |
| Mouse | Thoracic | 0.7 | 0.04 | 17.5 | 14.05 | 13.69 | 8.23 | 7.45 |
| Rat | Cervical | 0.01 | 0.3 | 0.033 | 0.47 | 0.30 | 0.08 | 0.11 |
| Rat | Cervical | 0.03 | 0.3 | 0.1 | 0.70 | 0.05 | 0.14 | 0.42 |
| Rat | Cervical | 0.1 | 0.3 | 0.333 | 0.19 | 1.13 | 1.28 | 1.40 |
| Rat | Cervical | 0.3 | 0.3 | 1 | 1.83 | 2.59 | 0.30 | 2.20 |
| Rat | Cervical | 1 | 0.3 | 3.333 | 6.19 | 8.18 | 7.26 | 7.36 |
| Rat | Cervical | 3 | 0.3 | 10 | 36.14 | 19.27 | 18.56 | N/A |
| Rat | Thoracic | 0.01 | 0.3 | 0.033 | 0.60 | 1.24 | 0.05 | 0.78 |
| Rat | Thoracic | 0.03 | 0.3 | 0.1 | 2.91 | 0.55 | 0.27 | 1.36 |
| Rat | Thoracic | 0.1 | 0.3 | 0.333 | 3.17 | 3.94 | 5.76 | 4.41 |
| Rat | Thoracic | 0.3 | 0.3 | 1 | 9.95 | 12.14 | 1.12 | 5.63 |
| Rat | Thoracic | 1 | 0.3 | 3.333 | 26.58 | 13.15 | 14.95 | 11.77 |
| Rat | Thoracic | 3 | 0.3 | 10 | 60.29 | 25.86 | 29.71 | N/A |
| Rat | Lumbar | 0.01 | 0.3 | 0.033 | 0.85 | 1.30 | 0.11 | 1.03 |
| Rat | Lumbar | 0.03 | 0.3 | 0.1 | 4.01 | 1.14 | 0.62 | 1.61 |
| Rat | Lumbar | 0.1 | 0.3 | 0.333 | 5.58 | 8.24 | 8.92 | 9.40 |
| Rat | Lumbar | 0.3 | 0.3 | 1 | 15.62 | 19.12 | 1.76 | 15.73 |
| Rat | Lumbar | 1 | 0.3 | 3.333 | 38.63 | 23.67 | 30.63 | 42.14 |
| Rat | Lumbar | 3 | 0.3 | 10 | 57.64 | 46.15 | 62.68 | N/A |
| NHP | Cervical | 75 | 15 | 5 | 2.11 | 14.99 | 20.77 | 10.71 |
| NHP | Thoracic | 75 | 15 | 5 | 2.60 | 18.56 | 30.90 | 19.71 |
| NHP | Lumbar | 75 | 15 | 5 | 5.88 | 23.22 | 49.87 | 24.42 |

N/A, Data not available due to technical issues

**Supplementary table 4.**
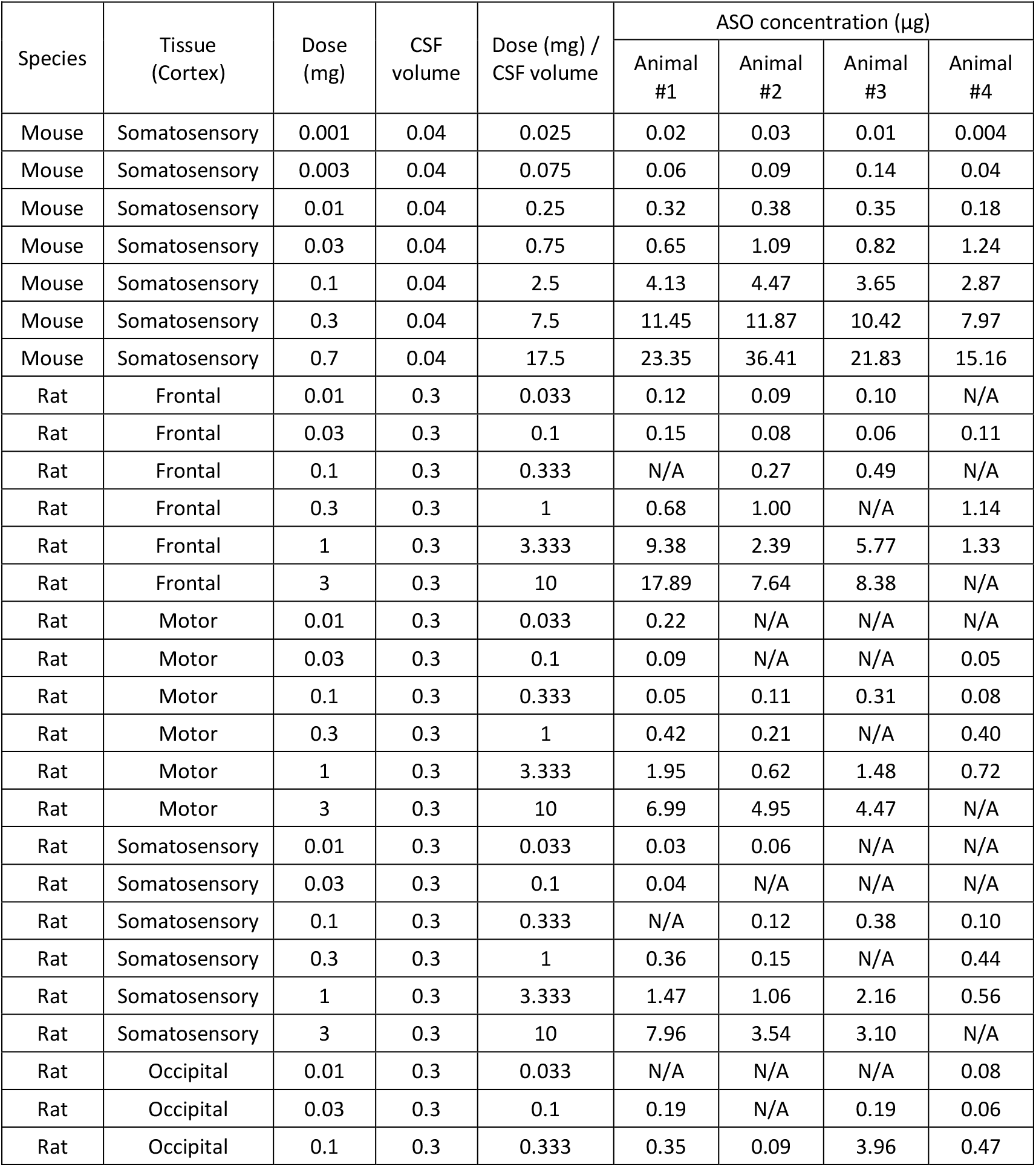
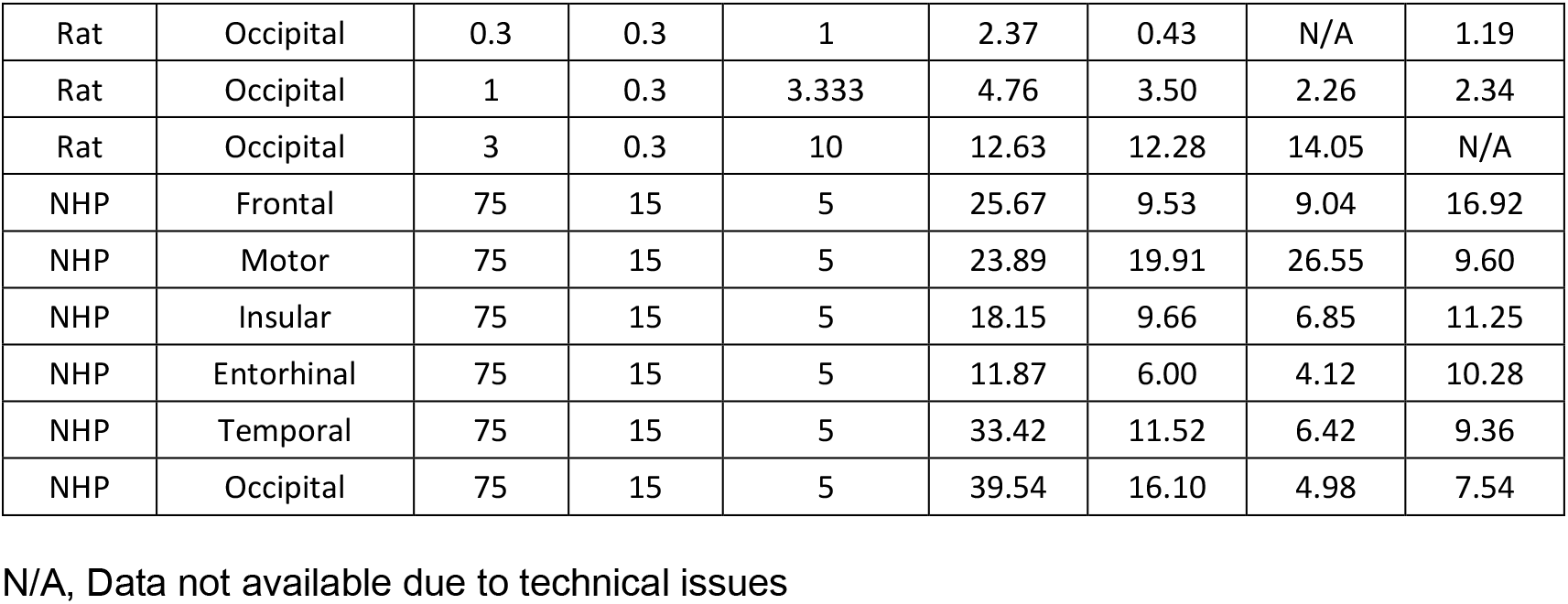
Cortex ASO tissue concentrations used in figure 10B.

| Species | Tissue (Cortex) | Dose (mg) | CSF volume | Dose (mg) / CSF volume | ASO concentration (µg) |  |  |  |
| --- | --- | --- | --- | --- | --- | --- | --- | --- |
|  |  |  |  |  | Animal #1 | Animal #2 | Animal #3 | Animal #4 |
| Mouse | Somatosensory | 0.001 | 0.04 | 0.025 | 0.02 | 0.03 | 0.01 | 0.004 |
| Mouse | Somatosensory | 0.003 | 0.04 | 0.075 | 0.06 | 0.09 | 0.14 | 0.04 |
| Mouse | Somatosensory | 0.01 | 0.04 | 0.25 | 0.32 | 0.38 | 0.35 | 0.18 |
| Mouse | Somatosensory | 0.03 | 0.04 | 0.75 | 0.65 | 1.09 | 0.82 | 1.24 |
| Mouse | Somatosensory | 0.1 | 0.04 | 2.5 | 4.13 | 4.47 | 3.65 | 2.87 |
| Mouse | Somatosensory | 0.3 | 0.04 | 7.5 | 11.45 | 11.87 | 10.42 | 7.97 |
| Mouse | Somatosensory | 0.7 | 0.04 | 17.5 | 23.35 | 36.41 | 21.83 | 15.16 |
| Rat | Frontal | 0.01 | 0.3 | 0.033 | 0.12 | 0.09 | 0.10 | N/A |
| Rat | Frontal | 0.03 | 0.3 | 0.1 | 0.15 | 0.08 | 0.06 | 0.11 |
| Rat | Frontal | 0.1 | 0.3 | 0.333 | N/A | 0.27 | 0.49 | N/A |
| Rat | Frontal | 0.3 | 0.3 | 1 | 0.68 | 1.00 | N/A | 1.14 |
| Rat | Frontal | 1 | 0.3 | 3.333 | 9.38 | 2.39 | 5.77 | 1.33 |
| Rat | Frontal | 3 | 0.3 | 10 | 17.89 | 7.64 | 8.38 | N/A |
| Rat | Motor | 0.01 | 0.3 | 0.033 | 0.22 | N/A | N/A | N/A |
| Rat | Motor | 0.03 | 0.3 | 0.1 | 0.09 | N/A | N/A | 0.05 |
| Rat | Motor | 0.1 | 0.3 | 0.333 | 0.05 | 0.11 | 0.31 | 0.08 |
| Rat | Motor | 0.3 | 0.3 | 1 | 0.42 | 0.21 | N/A | 0.40 |
| Rat | Motor | 1 | 0.3 | 3.333 | 1.95 | 0.62 | 1.48 | 0.72 |
| Rat | Motor | 3 | 0.3 | 10 | 6.99 | 4.95 | 4.47 | N/A |
| Rat | Somatosensory | 0.01 | 0.3 | 0.033 | 0.03 | 0.06 | N/A | N/A |
| Rat | Somatosensory | 0.03 | 0.3 | 0.1 | 0.04 | N/A | N/A | N/A |
| Rat | Somatosensory | 0.1 | 0.3 | 0.333 | N/A | 0.12 | 0.38 | 0.10 |
| Rat | Somatosensory | 0.3 | 0.3 | 1 | 0.36 | 0.15 | N/A | 0.44 |
| Rat | Somatosensory | 1 | 0.3 | 3.333 | 1.47 | 1.06 | 2.16 | 0.56 |
| Rat | Somatosensory | 3 | 0.3 | 10 | 7.96 | 3.54 | 3.10 | N/A |
| Rat | Occipital | 0.01 | 0.3 | 0.033 | N/A | N/A | N/A | 0.08 |
| Rat | Occipital | 0.03 | 0.3 | 0.1 | 0.19 | N/A | 0.19 | 0.06 |
| Rat | Occipital | 0.1 | 0.3 | 0.333 | 0.35 | 0.09 | 3.96 | 0.47 |
| Rat | Occipital | 0.3 | 0.3 | 1 | 2.37 | 0.43 | N/A | 1.19 |
| Rat | Occipital | 1 | 0.3 | 3.333 | 4.76 | 3.50 | 2.26 | 2.34 |
| Rat | Occipital | 3 | 0.3 | 10 | 12.63 | 12.28 | 14.05 | N/A |
| NHP | Frontal | 75 | 15 | 5 | 25.67 | 9.53 | 9.04 | 16.92 |
| NHP | Motor | 75 | 15 | 5 | 23.89 | 19.91 | 26.55 | 9.60 |
| NHP | Insular | 75 | 15 | 5 | 18.15 | 9.66 | 6.85 | 11.25 |
| NHP | Entorhinal | 75 | 15 | 5 | 11.87 | 6.00 | 4.12 | 10.28 |
| NHP | Temporal | 75 | 15 | 5 | 33.42 | 11.52 | 6.42 | 9.36 |
| NHP | Occipital | 75 | 15 | 5 | 39.54 | 16.10 | 4.98 | 7.54 |

N/A, Data not available due to technical issues

**Supplementary table 5.** Spinal cord ASO tissue concentrations for monkey simulated from rodent dose response used in figure 10A.

| Spinal cord ASO concentrations in monkey |  |
| --- | --- |
| Simulated based on Mouse dose response | Simulated based on Rat dose response |
| 2.99 | 17.88 |
| 2.98 | 13.25 |
| 3.19 | 23.67 |
| 2.98 | 17.08 |
| 3.15 | 17.75 |
| 3.08 | 19.69 |
| 2.86 | 24.08 |
| 2.82 | 21.51 |
| 2.95 | 20.20 |
| 2.82 | 13.30 |
| 2.94 | 22.81 |
| 2.93 | 19.35 |

**Supplementary table 6.** Cortex ASO tissue concentrations for monkey simulated from rodent dose response used in figure 10B.

| Cortex ASO concentrations in monkey |  |
| --- | --- |
| Simulated based on Mouse dose response | Simulated based on Rat dose response |
| 3.05 | 7.05 |
| 2.80 | 6.18 |
| 3.16 | 6.84 |
| 3.30 | 6.16 |
| 3.06 | 6.33 |
| 3.77 | 7.46 |
| 3.02 | 6.88 |
| 3.38 | 7.87 |
| 2.77 | 6.66 |
| 3.55 | 7.11 |
| 3.07 | 7.54 |
| 3.11 | 6.33 |
| 3.54 | 7.28 |
| 4.28 | 6.55 |
| 3.29 | 6.91 |
| 3.06 | 7.50 |
| 3.40 | 8.31 |
| 3.28 | 7.00 |
| 3.27 | 7.51 |
| 3.76 | 7.51 |
| 2.85 | 7.18 |
| 2.74 | 7.06 |
| 3.05 | 6.98 |
| 3.69 | 6.42 |

**Supplementary table 7.** Comparison of intercept and slope of mouse and rat regression lines in spinal cord and cortex

| Tissue | Species | Intercept |  |  |  | Slope |  |  |  |
| --- | --- | --- | --- | --- | --- | --- | --- | --- | --- |
|  |  | Mean | SEM | Difference | p-value | Mean | SEM | Difference | p-value |
| Spinal cord | Mouse | -0.237 | 0.0303 | -0.964 | <0.0001 | 1.015 | 0.031 | 0.194 | 0.0737 |
|  | Rat | 0.727 | 0.0588 |  |  | 0.821 | 0.067 |  |  |
| Cortex | Mouse | 0.0643 | 0.039 | 0.282 | 0.00153 | 1.112 | 0.0402 | 0.069 | 0.617 |
|  | Rat | -0.218 | 0.0574 |  |  | 1.043 | 0.0947 |  |  |

t-test comparisons - two-tailed

**Supplementary table 8.** Comparison of ASO tissue concentrations measured in monkey with predicted ASO tissue concentrations in mouse and rat at 5 mg/ml (Dose / CSF volume)

| Tissue | Comparison | DF | Difference | t Ratio | P value |
| --- | --- | --- | --- | --- | --- |
| Spinal cord | NHP v. mouse | 33 | 15.6693 | 4.81 | <0.0001 |
|  | NHP v. rat | 33 | -0.5694 | -0.17 | 0.8622 |
| Cortex | NHP v. mouse | 69 | 7.48098 | 4.79 | <0.0001 |
|  | NHP v. rat | 69 | 11.24651 | 7.2 | <0.0001 |

ANOVA, Multiple comparison test (t-test all pairwise)

## Notes

### Competing Interest Statement

The authors have declared no competing interest.

